# A Hybrid Cellular-Heterogeneous Catalyst Strategy for the Production of Olefins from Glucose

**DOI:** 10.1101/2020.11.17.387563

**Authors:** Zhen Q. Wang, Heng Song, Noritaka Hara, Dae Sung Park, Gaurav Kumar, Yejin Min, Paul J. Dauenhauer, Michelle C. Y. Chang

## Abstract

Living systems provide a promising approach to chemical synthesis, having been optimized by evolution to convert renewable carbon sources such as glucose to an enormous range of small molecules. However, a large number of synthetic structures can still be difficult to obtain solely from cells, such as unsubstituted hydrocarbons. In this work, we demonstrate the use of a hybrid cellular-heterogeneous catalytic strategy to produce olefins from glucose, using a selective hydrolase to generate an activated intermediate that is readily deoxygenated. Using a new family of iterative thiolase enzymes, we have genetically engineered a microbial strain that produces 4.3 ± 0.4 g L^−1^ of fatty acid from glucose with 86% captured as 3-hydroxyoctanoic and 3-hydroxydecanoic acids. This 3-hydroxy substituent serves as a leaving group enabling heterogeneous tandem decarboxylation-dehydration routes to olefinic products on Lewis acidic catalysts without the additional redox input required for enzymatic or chemical deoxygenation of simple fatty acids.

---

The ability to transform raw materials into functional fine and bulk chemicals has served as the basis for advances in modern society. As traditional chemical production has matured, new approaches to scalable catalysis have the potential to enable access to new chemical structures, take advantage of renewable resources, and lower environmental impact. In particular, living systems allow us to grow molecules from plant-sourced material in a single-stage fermentation at ambient temperature, pressure, and under aqueous conditions.^1–5^ Despite the enormous breadth of molecular structures found in Nature that can be produced from glucose, there still remains a large range of small molecules made through synthetic chemistry that are not readily accessed through typical enzymatic pathways.

One group of these molecules are those originating from petroleum feedstocks, which tend to provide highly reduced and unmodified hydrocarbons used as fuels, lubricants, and polymer precursors (**Fig. 1A**). In comparison, their biologically-derived counterparts tend to be highly functionalized, consisting mostly of esters and alcohols rather than true hydrocarbons. Enzymes involved in wax synthesis or secondary metabolism have been found to decarboxylate or decarbonylate unsubstituted fatty acids and aldehydes to generate hydrocarbon structures.^6–12^ However, these require Fe and O_2_ redox chemistry while also mostly working with either long-chain substrates (<C_18_) or complex biosynthetic intermediates.

**Figure 1.**
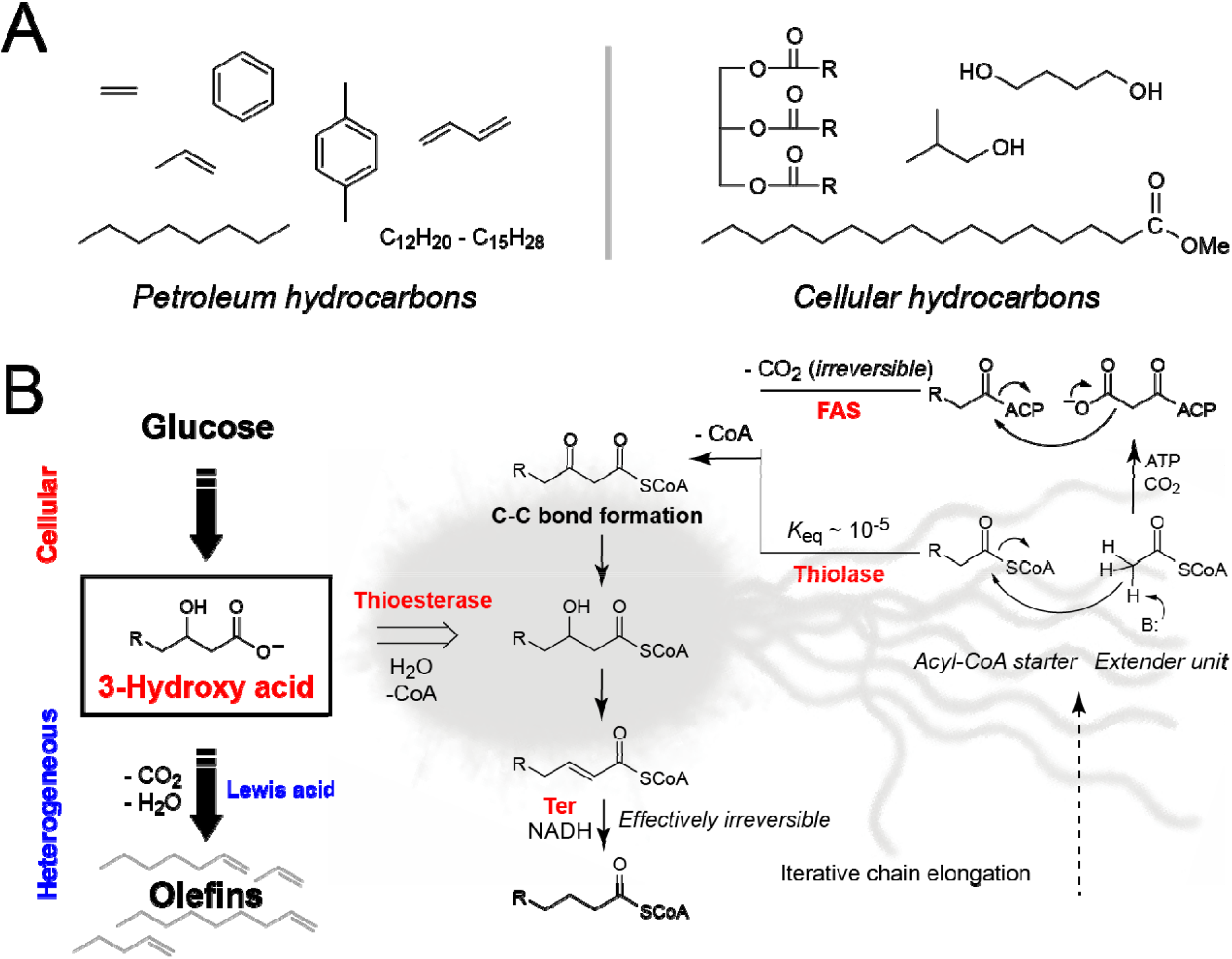
Biological hydrocarbons. (A) Structures of petroleum-derived hydrocarbons compared to those typically made in living systems. (R, naturally-occurring acyl chains are typically C_16_-C_20_ in length and contain 0-3 double bonds) (B) The design of a FAS-independent pathway based on iterative thiolases where the malonyl-CoA extender unit is replaced by acetyl-CoA. FASs utilize malonyl-CoA extender units attached to an ACP carrier to drive the C-C bond formation equilibrium by the irreversible loss of CO_2_. The biosynthesis of these extender units requires ATP and is highly regulated inside cells. While thiolases can carry out a Claisen condensation between two unactivated acyl-CoA substrates, the reaction equilibrium strongly favors bond cleavage rather than bond formation. However, the use of the effectively irreversible enoyl-CoA reduction catalyzed by *trans*-enoyl-CoA reductase (Ter) enzymes can replace this driving force in order to draw each chain elongation cycle forward in a downstream step. A thioesterase selects both the chain length and oxidation state of the acyl-CoA to exit the cycle, producing a free fatty acid. Use of a medium chain 3-hydroxy acid-selective thioesterase produces the 3-hydroxyoctanoic and 3-hydroxydecanoic acids, which are activated by solid Lewis acid catalysts for tandem decarboxylation-dehydration routes via loss of water and carbon dioxide, resulting in formation of olefin products without the need for redox chemistry. (ACP, acyl carrier protein)

In this work, we report the design and engineering of a fatty acid synthase (FAS)-independent pathway for the microbial production of medium-chain 3-hydroxy acids. To construct this pathway, we identified and characterized a new family of thiolases, which can carry out the iterative chain elongation of acetyl-coenzyme A (CoA) extender units (**Fig. 1B**). Although this approach lifts the high ATP cost of using activated malonyl-CoA extender units, the driving force for carboncarbon bond formation from coupling to the release of carbon dioxide is also lost. To compensate for the low *K*_eq_ of the thiolase reaction (10^−5^), we instead use the downstream *trans-enoyl* reductase (Ter) to draw the pathway equilibrium forward.^13,14^

To exit the elongation cycle, we use a thioesterase that selectively produces C_8_ and C_10_ 3-hydroxy acids, which can be utilized as precursors for the corresponding (*n*-1) terminal alkene **(Fig. 1B**). Enzymatic processing of 3-hydroxy acids requires activation of the leaving group through ATP-dependent phosphorylation or PAPS-dependent sulfation while also being highly restricted in substrate scope.^10,11^ As such, we pair cellular production of 3-hydroxy acids with a downstream heterogeneous process to produce a range of short- and medium-size olefinic products using redox-free Lewis acid catalysis. These olefins are particularly useful as the flexible alkyl branches of medium chain-length can yield viscous liquids even at low temperature when incorporated into polymer products. In addition, alkenes can be used for a variety of downstream chemical processes including hydrogenation, hydration, halogenation, and epoxidation. This hybrid approach serves as a general platform that takes advantage of both the ability of living systems to utilize renewable feedstocks for scalable synthesis as well as the flexibility of heterogeneous catalysis for rapid processing to form valuable structures.

## Results and discussion

### Discovery and characterization of a new family of iterative thiolases for fatty acid biosynthesis

For the most part, hydrocarbons are produced in living systems to serve as membrane and storage lipids and rely on the iterative chain elongation catalyzed by FAS enzymes. Accordingly, FASs can support high production levels of lipid.^15–17^ However, this process is also highly regulated and energy intensive.^18^ More specifically, FAS pathways are tightly coupled to cell growth phase and subject to feedback control and oxygen availability. They also require stoichiometric protein carriers (acyl carrier protein, ACP) and ATP used for the activation of acetyl-CoA to generate the malonyl-CoA extender unit (**Fig. 1B**). Given these intrinsic limitations, we sought to identify a FAS-independent system for the biosynthesis of medium-chain hydrocarbons that could be decoupled from its strict metabolic regulation and dependence on ATP, ACP, and O_2_.

These requirements can be fulfilled by thiolases, which carry out the Claisen condensation of two acyl-CoA substrates to produce a 3-ketoacyl-CoA intermediate similar to that found in FASs. Indeed, the promiscuous activity of the two known thiolases, BktB and FadA, can be run in retro-degradative direction to produce a variety of medium-chain alcohols and carboxylic acids^19–21^ Despite their demonstrated utility in biosynthesis and the large number of family members (>50,000), only a handful of thiolases have been characterized with regard to native substrate selectivity and have mostly been found to carry out dimerization to form short-chain products or degradation of long-chain substrates.^22,23^ We therefore set out to identify new biosynthetic thiolases distinct from those participating in polyhydroxyalkanoate (PHA) synthesis or fatty acid degradation (β-oxidation) with the overall goal of discovering enzymes with new biochemical activities.

Towards this goal, thiolase sequences in the Pfam database were reduced to representative sequences below 60% identity for transitive clustering using Cytoscape.^24,25^ As expected, most of the thiolases were clustered into the two major nodes represented by short-chain thiolases involved in PHA synthesis (PhaA/BktB, C_2_-C_6_)^26,27^ or broadrange thiolases used for fatty acid degradation (FadA, C_2_-C_16_)^23^ (**Fig. 2A**). PaaJ thiolases^28^ involved in the bacterial phenylacetate catabolic pathway were located between the PhaA and FadA nodes. Interestingly, we found a branch of PaaJ-like thiolases from Mycobacteria that was well separated from other known thiolases. When examining the genomic context of these PaaJ-like thiolases, we noted that these thiolases were separate from the genes for β-oxidation and also that several were clustered with genes annotated as 3-ketoacyl reductases (*fabG4*), enoylacyl hydratases (*maoC*), and acyl dehydrogenases (*fadE* or *caiA*) (**Supplemental Table S1**, **Supplemental Fig. S1A**)^29–32^ This finding suggested to us that these PaaJ-like thiolases could participate in an anabolic pathway for CoA-dependent hydrocarbon elongation system orthogonal to native fatty acid synthesis (**Fig. 2B**).

**Figure 2.**
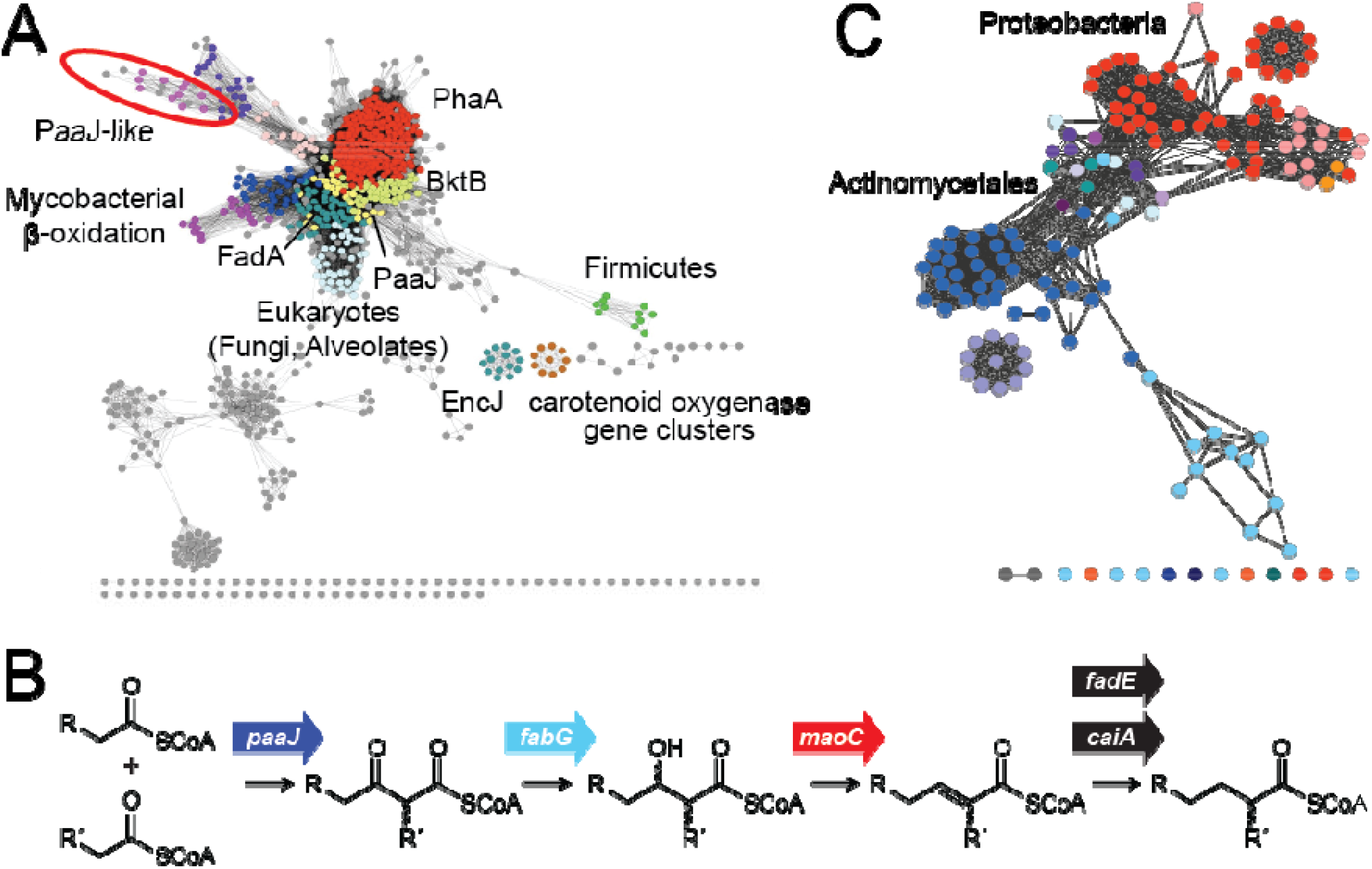
Identification of a new biosynthetic PaaJ-like thiolase family. (A) Transitive clustering of representative thiolases. The PaaJ-like branch from mycobacteria is highlighted with a red oval.(PhaA, acetyl-CoA acetyltransferase from polyhydroxybutyrate (PHB) biosynthesis; BktB, β-ketothiolase from polyhydroxyvalerate (PHV) biosynthesis; PaaJ, 3-ketoacyl-CoA thiolase from phenylacetic acid catabolism; FadA, 3-ketoacyl-CoA thiolase from fatty acid β-oxidation; EncJ: β-ketoacyl-CoA thiolase in benzoyl-CoA biosynthesis. (B) Fatty acid catabolic genes found in the mycobacterial PaaJ-like clusters. The thiolase (PaaJ) catalyzes Claisen condensation between two acyl-CoA substrates. Ketoreduction can be catalyzed by a 3-keto-acyl-CoA reductase (FabG), followed by dehydration catalyzed by a enoylacyl-CoA hydratase (MaoC). Enoyl reduction catalyzed by an enoylacyl-CoA reductase (CaiA or FadE) produces the fully reduced acyl-CoA species that can undergo another round of chain elongation or other downstream processing reactions.

Since *Ms*PaaJ1 from *Mycobacterium smegmatis* (**Supplemental Table S1**) demonstrated issues with solubility, we expanded the PaaJ-like node to search for orthologs with potentially improved solubility (**Fig. 2C**). This expanded network revealed additional PaaJ-like thiolases in the Proteobacteria phylum to which *E. coli* belongs. From these sequences, we selected nine (PaaJ1-9) that were also clustered with genes annotated with fatty acid anabolic activities while also being distinct from β-oxidation clusters. These gene sequences were codon optimized for expression in *E. coli* and the proteins found to be mostly expressed in soluble form (**Supplemental Table S1**, **Supplemental Fig. S1B**).

PaaJ1-9 were first screened for their ability to catalyze CoA release with acetyl-CoA (C_2_) and butyryl-CoA (C_4_) (**Supplemental Fig. S2**). Further experiments with the four active thiolases with longer-chain linear acyl-CoA starters (C_2_-C_10_) revealed that PaaJ5, PaaJ7, and PaaJ9 appeared to be capable of iterative condensation (**Supplemental Fig. S3**). While all of the three thiolases could accept short- and medium-chain linear acyl-CoAs as substrates in the condensing direction, they appear to favor hexanoyl-CoA over acetyl-CoA suggesting that these enzymes comprise a new family of thiolases selective for medium-chain acyl-CoAs.

### Design and *in vitro* reconstitution of a pathway for thiolase-dependent medium-chain fatty acid synthesis

In addition to the initial carbon-carbon bond forming step catalyzed by thiolases, three additional steps of ketoreduction, dehydration, and enoyl reduction are needed to close the elongation cycle (**Fig. 3A**). A thioesterase is also needed to select the intermediate of the desired chain length to generate the free fatty acid product and recycle the CoA cofactor. To construct this pathway, we utilized genes derived from these PaaJ-like thiolase gene clusters and their orthologs as well as the *trans*-enoyl reductase from *Treponema denticola* (*Td*Ter) to provide an effectively irreversible step to drive the elongation cycle forward.^13,14^

**Figure 3.**
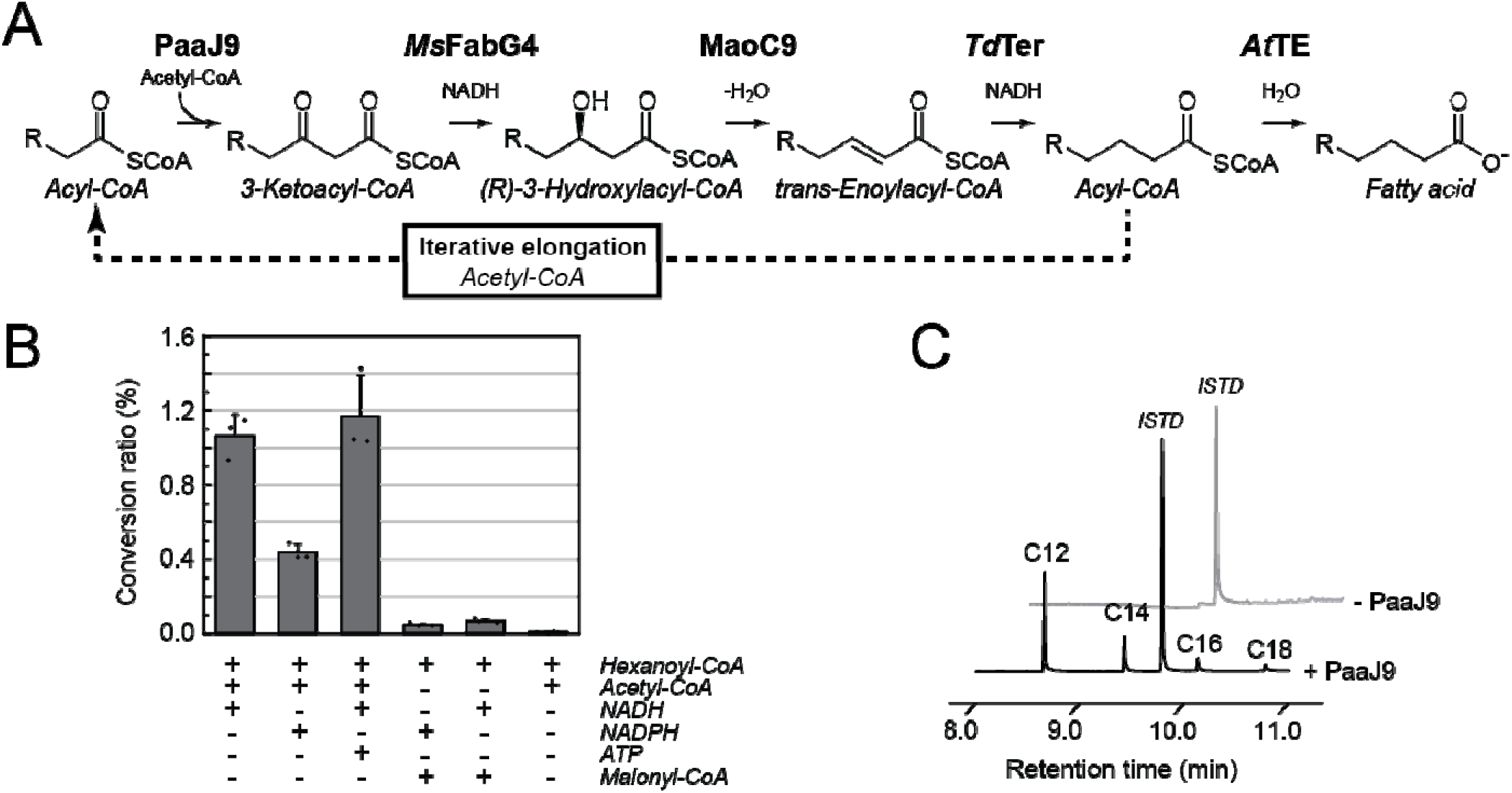
*In vitro* reconstitution of the PaaJ-dependent fatty acid synthesis pathway. (A) Enzymes and reactions used to reconstitute the pathway (PaaJ9, PaaJ-like thiolase from *Pseudomonas putida; Ms*FabG4, 3-keto reductase from *M. smegmatis;* MaoC9, enoylacyl-CoA reductase from *P. putida; Td*Ter: *trans*-enoylacyl-CoA reductase from *T. denticola; *At*TE:* acyl-CoA thioesterase from *A. tetradium*) (B) Cofactor requirements of the thiolase-dependent carbon single-step elongation system containing purified PaaJ9, *Ms*FabG4, MaoC9, *Td*Ter, and *At*TE. Cofactors and substrates were added as noted. The conversion ratio was calculated as the molar percentage of hexadecanoic acid converted to octadecanoic acid. Data are mean ± s.d. of technical replicates (n = 3). (C) *In vitro* iterative elongation with acetyl-CoA and NADH using the four-enzyme system consisting of purified PaaJ9, *Ms*FabG4, MaoC9, and *Td*Ter in the absence of *At*TE. The reaction was analyzed by GC-MS after extraction and functionalization of the fatty acid products. The chromatogram shown is representative of 3 biological replicates (C_12_, dodecanoic acid; C_14_, tetradecanoic acid; C_16_, hexadecanoic acid; C_18_, octadecanoic acid).

We selected two 3-ketoacyl-CoA reductases to characterize, *Ms*FabG4 and FabG9 from the *Ms*PaaJ1 and PaaJ9 clusters, respectively (**Supplemental Table S1**, **Supplemental Fig. S1**). While purified FabG9 was not found to catalyze reduction of acetoacetyl-CoA, MsFabG4 was determined to be a NADH-dependent *(R)*-3-ketoacyl-CoA reductase with a preference for C_6_ substrates over C_4_ (**Supplemental Fig. S4**). Therefore, *Ms*FabG4 demonstrates significant differences from canonical FabGs, which utilize acyl-ACP substrates and NADPH. *Ms*FabG4 also differs from the FadB enzymes of the fatty acid *β-* oxidation because the latter are specific to *(S)*-3-hydroxylacyl-CoAs.^23^

For the next step, the 3-hydroxylacyl-CoA dehydratase from the PaaJ9 cluster, MaoC9, was characterized as a *(R)*-3-hydroxyacyl-CoA dehydratase and therefore stereochemically compatible with FabG4 (**Supplemental Fig. S4**). To close the cycle, *Td*Ter was selected both for its ability to drive thiolase-dependent pathway equilibrium forward as well as its broad substrate selectivity, demonstrating high activity on linear C_4_, C_6_, and C_12_ substrates.^33^

With a full set of genes for iterative chain elongation in place, we turned our attention to identification of a thioesterase to produce the desired C_8_-C_10_ products. We selected several thioesterases from the literature that were reported to demonstrate selectivity towards medium-chain acyl-ACP substrates,^34,35^ however, *in vivo* experiments suggested that these thioesterases were not effective on medium-chain acyl-CoA substrates (**Supplemental Fig. S5A**). We then tested a thioesterase candidate from *Anaerococcus tetradium* (*At*TE), which was reported to hydrolyze octanoyl-ACP but was not phylogenetically grouped with other known thioesterases^36^ *In vitro* characterization of purified *At*TE demonstrated that this enzyme selectively hydrolyzed medium-chain acyl-CoAs (C_8_-C_10_) while little activity was detected for longer or shorter-chain acyl-CoAs (**Supplemental Fig. S5B**).

A five-enzyme system consisting of purified PaaJ9, *Ms*FabG4, MaoC9, *Td*Ter, and *At*TE was reconstituted with hexanoyl-CoA, as a starter, along with acetyl-CoA and NADH. End product analysis showed that the expected octanoic acid was made along with trace levels of decanoic acid, 3-hydroxyoctanoic acid, and 3-hydroxydecanoic acid, suggesting that *At*TE may also hydrolyze pathway intermediates such as the C_8_ and C_10_ 3-hydroxyacyl-CoAs (**Supplemental Fig. S6A**). The cofactor requirements were also studied by measuring the conversion ratio of hexanoyl-CoA to octanoic acid, demonstrating that it was ATP- and malonyl-CoA-independent as expected and that NADH is its preferred cofactor over NADPH (**Fig. 3B**).

In order to examine iterative chain elongation without an added starter unit, *At*TE was excluded from the assay to avoid causing premature release of intermediates. With only acetyl-CoA and NADH as substrates, this four-enzyme system produced dodecanoyl-, tetradecanoyl-, hexadecanoyl-, and octadecanoyl-CoAs with dodecanoyl-CoA as the dominant product (**Fig. 3C**. Interestingly, we do not find smaller fatty acid products, perhaps because the matched selectivities of the enzymes reduce the level of early exit of intermediates from the elongation cycle (**Supporting Fig. S6B**). Based on its higher *in vitro* activity, PaaJ7 was also tested in the reconstituted system but demonstrated almost 5-fold lower productivity compared to PaaJ9 (**Supplemental Fig. S6C**). Based on these data, the designed PaaJ9 thiolase pathway is capable of completing 5-8 cycles of iterative carbon elongation from an acetyl-CoA starter with an apparent chain length preference of C_12_.

### Construction of a pathway for medium-chain 3-hydroxy fatty acid synthesis in *E. coli*

Having validated the *in vitro* activities of the pathway enzymes, we set out to construct a synthetic pathway for medium-chain fatty acid production in a microbial host. We developed an initial two-plasmid system where the genes encoding PaaJ9, FabG9, and MaoC9 were cloned into a high-copy pCDF2 backbone plasmid behind a double *tac* promoter with the hypothesis that genes derived from the same native cluster could potentially operate best together *in vivo*. The remaining genes encoding *Td*Ter and a thioesterase (TesB, *Cp*FatB1, or *Af*TE) were then inserted into a second pCWori-based plasmid (**Fig. 4A**). These two plasmids were transformed into *E. coli* DH1(DE3) and the resulting strains were tested for their abilities to produce medium-chain fatty acids under microaerobic conditions (**Fig. 4B**, strains **1**-**3**). Analysis of the product profile showed that it was dominated by octanoic and decanoic acids and that the overall titers were dependent on the thioesterase. The highest titer was obtained with *At*TE (strain **3**), which produced a total of 134 ± 17 mg L^−1^ octanoic acid, 3-hydroxyoctanoic acid and 3-hydroxydecanoic acid.

**Figure 4.**
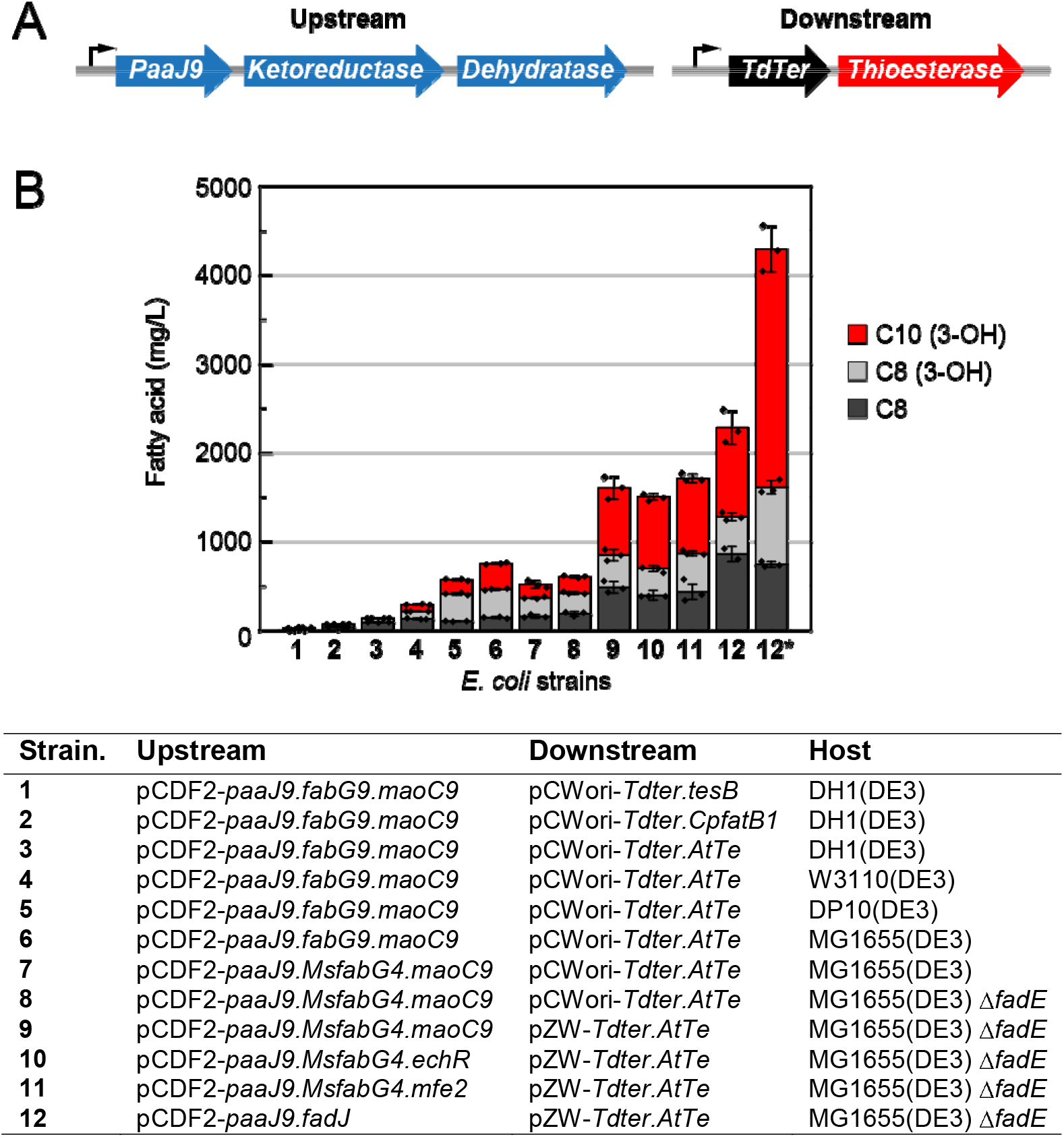
Production of medium-chain C_8_-C_10_ fatty acids in engineered E. coli strains containing a thiolase-dependent pathway for iterative chain elongation. (A) Schematic diagram of upstream and downstream plasmids. (B) Fatty acid production in engineered strains. E. coli strains were co-transformed with both upstream and downstream plasmids and cultured for 3 d under microaerobic conditions at 30 °C in TB broth containing 2.5% (*w/v*) glucose. Strain **12\*** represents Strain 12 cultivated in a one-batch fermentation process in a 1 L bioreactor (C8, octanoic acid; C8 (3-OH), 3-hydroxyoctanoic acid, C10 (3-OH), 3-hydroxydecanoic acid). paaJ9, thiolase from P. putida; fabG9, 3-ketoacyl reductase from *P. putida*; *maoC9*, enoylacyl hydratase from *P. putida*; *Tdter, trans*-enoyl-CoA reductase from *T. denticola*; *tesB*, acyl-ACP thioesterase II from *E. coli*; *CpfatB1*, acyl-ACP thioesterase from *Cuphea palustris*; *AtTe*, acyl-CoA/ACP thioesterase from *A. tetradium*; *MsfabG4*, 3-ketoreductase from *M. smegmatis*; *echR*, enoyl-CoA hydratase from *Rickettsia prowazekii*; *mfe2*, enoyl-CoA hydratase 2 from Arabidopsis *thaliana*; *fadJ*, bi-functional 3-ketoacyl reductase/enoylacyl-CoA hydratase from *E. coli*; Data are mean ± s.d. of biological replicates (n = 3).

With a working pathway in hand, we screened various host strains to optimize production, leading to an overall 6-fold increase in C_8_-C_10_ acids to 780 ± 60 mg L^-1^ in MG1655(DE3) (**Fig. 4B**, strains **3–6**). Replacing FabG9 with its homolog *Ms*FabG4, the NADH-dependent reductase producing *(R)*-3-ketoacyl-CoAs, led to a higher ratio of octanoic acid over 3-hydroxy fatty acids but no increase in overall productivity (**Fig. 4B**, strain **7**). Further host optimization by knocking out *fadE*^17^ encoding the native acyl-CoA dehydrogenase in the β-oxidation pathway slightly increased overall yields presumably because of loss of the host ability to degrade intermediates (**Fig, 4B**, strain **8**).

Since high thioesterase expression can often lead to cell growth defects, we next transitioned to a **w**eak promoter for thioesterase expression, resulting in a 2-fold improvement of yield to 1600 ± 200 m**g** L^−1^. (**Fig. 4B**, strain **9**). Further analysis of strain **9 us**ing cell lysate assays indicated that *Ms*FabG4 and MaoC were bottlenecks (**Supplemental Fig. S7A**). Given that the production of 3-hydroxyacids could either arise from thioesterase promiscuity or a dehydratase limitation, we first screened homologs of the M**ao**C9 dehydratase (**Fig 4B**, strains **10** and **11**). However, the resulting strains did not provide higher ti**ter**s or changed ratios of 3-hydroxy fatty acids to octanoate. We therefore replaced *Ms*FabG4 and MaoC with the bifunctional ketoreductase-dehydratase FadJ from *E. coli*^37^. This replacement resulted in a 1.5-fold improvement in yield to 2210 ± 320 mg L^−1^ under microaerobic conditions (**Fig. 4B**, strain **12**).

Interestingly, studies of strain **12** suggest that eliminating the upstream thiolase pathway leads to reduced fitness and that the acyl-CoAs produced may provide sufficient substrate to circumvent thioesterase toxicity (**Supplemental Fig. S7B**). One possible interpretation of this data is that *At*TE does prefer acyl-CoA substrates to the acyl-ACP substrates generated by FAS-dependent pathways when both are available. Indeed, experiments comparing strains containing the full pathway to the thioesterase alone show that production drops to background levels and that *Af*TE appears to more effective on acyl-CoA substrates in this system (**Supplemental Fig. S5A** and **S7C**). Furthermore, studies with the FAS inhibitor, triclosan,^38^ demonstrate that the PaaJ9 thiolase pathway is indeed orthogonal to the host FAS system (**Supplemental Fig. S7D**).

Strain **12** was then grown in a 1-L bioreactor to better assess production (**Fig. 4B**, strain **12\***). During the one-batch fermentation process, a total hydrocarbon titer of 4.3 ± 0.4 g L^−1^ was achieved, with 3-hydroxy acids as the dominant product 3.6 ± 0.3 g L^−1^ and the yield of octanoic acid (0.74 ± 0.03 g L^−1^) approaching the maximum octanoate solubility in water.

### The semisynthesis of olefins from C_8_-C_10_ 3-hydroxy acids using heterogenous catalysis

The production of 3-hydroxy acids leads to the potential for different downstream chemistry compared to that of their simple fatty acid counterparts. Fatty acids and aldehydes can be decarboxylated or decarbonylated enzymatically to the (*n*-1) terminal alkene or alkane, respectively, through iron/O_2_-dependent radical processes^7−9,12^ Unsubstituted fatty acids can also be converted to their (n-1) alkane congeners by heterogeneous methods but this is also a relatively energetically intensive process.^17,39^ In the case of 3-hydroxy acids, they are already preactivated for decarboxylation without additional energy input as the hydroxyl group can serve as a leaving group to produce the (*n*-1) terminal alkene (**Fig. 5A**). Enzymes such as OleBC or the polyketide synthase CurM are capable of catalyzing this reaction but still require ATP or PAPS as a phosphoryl or sulfuryl donor to further activate the leaving group.^10,11^ Given the limited substrate scope of these enzymes, heterogeneous catalysis may provide a more general approach to develop this reaction and should more easily accommodate change in substrate structure as long as the 3-hydroxy acid moiety remains intact.

**Figure 5.**
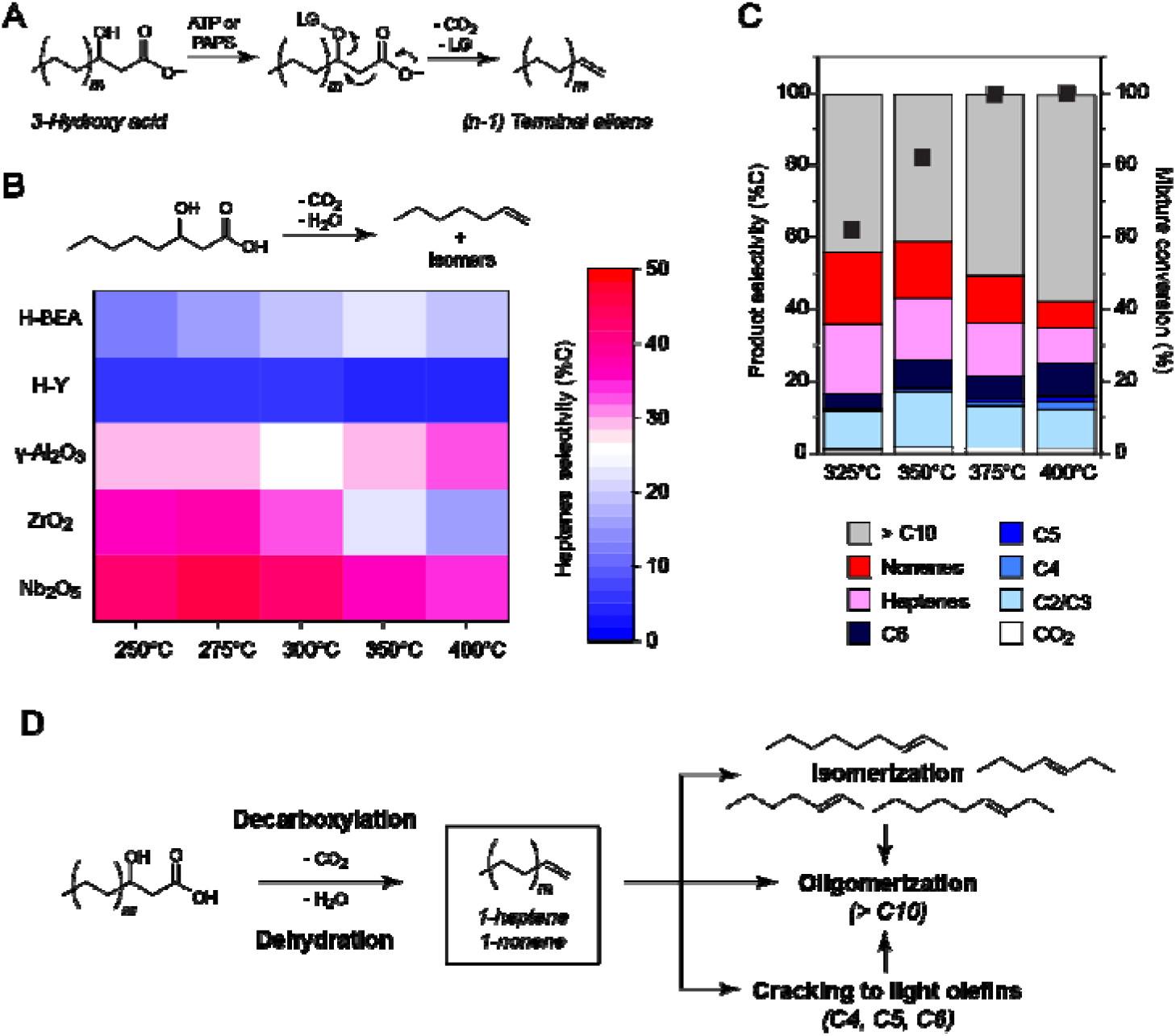
Downstream processing of 3-hydroxy fatty acids to olefins on heterogeneous catalysts. (A) The enzymatic decarboxylation of 3-hydroxy acid is initiated by activating the 3-hydroxy substituent as a leaving group (LG) for decarboxyation. (B) Screening on catalysts for tandem decarboxylation-dehydration of a model substrate. The heatmap depicts selectivity for production of heptenes from 3-hydroxyoctanoic acid on a variety of Brønsted and Lewis acidic catalysts at quantitative conversion of substrate. (C) Fermentation extracts from strain **12** containing 3-hydroxyoctanoic (*m* =2) and 3-hydroxydecanoic (*m* = 3) acids were converted to olefins on Nb2O5. The bar graph represents the olefin product distr**ibu**tion (*left axis*) while the black squares correspond to the conversion of the fermentation extract calculated from the sum of 3-hydroxoctanoic and 3-hydroxydecanoic acid peak areas. (Space velocity = 45 s^−1^) (D) Proposed reaction pathways where initial formation of 1-heptene (*m* = 2) and 1-nonene (m = 3) is followed by isomerization of the double bond to form heptenes and nonenes, cracking to light olefins, as well as oligomerization to larger species.

A combination of different Brønsted acidic large-pore-size zeolites (HY, H-BEA) as well as Lewis acid catalysts (ZrO_2_, γ-Al_2_O_3_, Nb_2_O_5_) were investigated for the tandem decarboxylation-dehydration of a 3-hydroxyoctanoic acid standard **(Fig. 5B**, **Supplemental Fig. S8**). In agreement with the ability of Brønsted acidic sites to catalyze oligomerization pathways,^40,41^ observed heptene selectivities were low (5-20%) on both H-BEA and H-Y. Alternatively, Lewis acidic catalysts exhibited higher heptene selectivities (30-45%), with niobium(V) oxide (Nb_2_O_5_) demonstrating highest propensity towards their formation (**Fig. 5B**). When applied to the microbial products extracted from fermentation broth, containing both 3-hydroxyoctanoic and 3-hydroxydecanoic acids, heptenes and nonenes were both found present in the reactor effluent with Nb_2_O_5_, with combined selectivities in the range 30-40% at moderate temperatures (<350 °C) (**Fig. 5C**). The product distribution on these solid acid catalysts is consistent with a reaction pathway where carbon dioxide and water are first eliminated from the 3-hydroxy acid to produce the corresponding (*n*-1) terminal alkene via a tandem decarboxylation-dehydration pathways (**Fig. 5D**).^42,43^ This terminal alkene can then either be isomerized to form internal alkenes, further cracked to smaller alkene products, or undergo complex condensation to heavy oligomerization products. Taken together, this hybrid approach to hydrocarbon production serves as a proof-of-concept platform to demonstrate the production of olefins from glucose using both cellular and heterogeneous catalysis.

## Conclusions

In conclusion, we have described a hybrid catalytic process that couples microbial fermentation to heterogeneous catalysis, enabling the production of olefins from glucose. This approach takes advantage of the ability of living organisms to convert simple renewable carbon feedstocks such as glucose to a broad range of structures, including functionalized hydrocarbons. However, the tailoring of these products using enzymatic chemistry can be challenging to implement with extensive protein engineering efforts. We therefore turned to heterogenous methods for deoxygenation to generate a hydrocarbon product.

Towards this goal, we have identified and characterized a new system of PaaJ-like thiolases that are capable of iterative chain elongation from the C_2_ starter, acetyl-CoA, to produce linear fatty acid products of C_12_-C_18_. This process was shown to be orthogonal to the native FAS system, overcoming its intrinsic limitations related to high ATP cost, O_2_- and ACP-dependence, as well as the tight cellular regulation of the malonyl-CoA pool and membrane homeostasis. However, the use of malonyl-CoA by FAS also provides an irreversible step for carbon-carbon bond formation that ultimately drives each chain elongation cycle forward. However, we have shown that this step can be substituted with an effectively irreversible enoyl reduction step downstream, which is sufficient to accumulate high levels of product. Optimization of this pathway *in vivo* yields microbial strains producing 4.3 ± 0.4 g L^−1^ of medium-chain (C_8_-C_10_) fatty acid product with the bulk produced as 3-hydroxy acid (86%).

By using thioesterase selectivity to provide an activating group for the fatty acid in the form of the 3-hydroxy substituent, we are able to redu**ce** the overall energetic cost for deoxygenation compared to enzymatic or chemical methods. Using heterogeneous catalysts, we have shown that it is possible to use simple Lewis acid catalysis in order to generate the deoxygenated product in the form of unfunctionalized olefins that can favor different size regimes. This hybrid process could potentially be applied to both linear and branched targets and lays the foundation for the production of bio-naphtha with a wide range of potential properties and applications.

## Methods

### *In vitro* reconstitution of the PaaJ-dependent pathway

*In vitro* pathway reconstitution was carried out in 300 μL reactions in 50 mM HEPES at pH 7.0 with 5 mM MgCl_2_ and 1 mM dithiothreitol (DTT) at 37 °C. For non-iterative reactions, purified recombinant PaaJ9 (2.5 μM), *Ms*FabG4 (1 μM), MaoC9 (1 μM), *Td*Ter (1 μM) and *At*TE (10 μM), and acetyl-CoA (5 mM). Reactions were initiated by adding hexanoyl-CoA (5 mM) and NADH (15 mM). When used, malonyl-CoA (5 mM), ATP (5 mM), and NADPH (15 mM) were also included. After 1 h, samples were quenched by addition of 6N HCl (100 μL). For iterative reactions, *At*TE was omitted from the reaction mixture to produce acyl-CoAs instead of free fatty acids. The hexanoyl-CoA starter was also omitted and acetyl-CoA (10 mM) was used to initiate the reaction. The acyl-CoA products were first hydrolyzed to free fatty acids by addition of 10 M sodium hydroxide (20 μL) and incubation at 65 °C for 20 min before acidification. The resulting samples were extracted with diethyl ether and derivatized with TMS-DAM for quantification against a standard curve by GC-MS (Thermo Scientific Trace GC Ultra-DSQII) after normalization with an internal standard.

### *In vivo* production of fatty acids

The microaerobic production of fatty acids in *E. coli* was performed based on literature methods.^13^ Briefly, chemically-competent *E. coli* strains were transformed with plasmids by heat shock. A single colony from freshly transformed plate was inoculated into Terrific Broth (TB) replacing the standard glycerol supplement with 1.5% (*w*/*v*) glucose and with appropriate antibiotics at 37 °C in a rotary shaker (200 rpm). Overnight cultures were inoculated into 25 mL TB with 1.5% (*w*/*v*) glucose and antibiotics in a 250 ml baffled flask at an optical density of 600 nm (OD_600_) at 0.05 in a rotary shaker (200 rpm) at 37 °C. When the OD_600_ reached 0.35 to 0.45, the culture was induced with 1.0 mM isopropyl β-D-1-thiogalactopyranoside (IPTG). The growth temperature was then reduced to 30 °C. The flasks were sealed with Parafilm M to prevent loss of volatile compounds produced and to limit oxygen in the flask. After 24 h, an additional 1% (*w*/*v*) glucose was added to the flasks and sealed again with Parafilm M. Samples were collected after 3 d for fatty acid analysis. Fatty acids were quantified by GC-MS as described. 3-Hydroxy acids were quantified against a standard curve by LC-MS using an Agilent 1290 HPLC coupled to a 6460 triple-quadrupole MS in negative ion mode and normalized with an internal standard.

### Fermentation

For larger-scale production of medium-chain fatty acids, one-batch cultures were carried out in a fermentor (Sartorius Stedim Biotech GmbH, Germany) containing 1 L of terrific broth medium supplemented with 5% (*w*/*v*) glucose as the sole carbon source and antibiotics (100 ug/mL carbenicillin, and 100 ug/mL spectinomycin). The seed cultures were incubated overnight at 37 °C and 250 rpm to use for sub-culturing into the same medium (50 mL). The sub-culture was inoculated to an OD_600_ of 1.0 and grown at 37 °C and 250 rpm to an OD_600_ of 3–5 (~ 3 h). These cells were used to inoculate a 1 L bioreactor at 5% (*v*/*v*) containing the same medium and 1 mM IPTG. Cultivation was carried out at 30 °C with the pH was adjusted to 7.0 using sodium hydroxide. The airflow and the initial agitation rate were set at 2 VVM and 700 rpm, respectively. The dissolved oxygen tension was controlled at 80% of saturation using an agitation cascade. The fermentation was allowed to proceed for 2 d before sample analysis and product extraction.

### Conversion of 3-hydroxy fatty acids to olefins

The fermentation supernatant (500 mL) was treated with conc. HCl to adjust the pH to 1.0 and extracted with 3 × 200 mL diethyl ether. The organic layers were combined, dried over MgSO_4_, and evaporated to dryness by rotary evaporation to provide the fatty acid extracts. The catalytic conversion of all fatty acids extracts were carried out using a pulsed flow microreactor integrated with a gas-chromatograph (GC) highlighted as previously described.^39^ Briefly, the fatty acids were dosed from the autosampler of a GC (Agilent 7890A) as pulses to the front inlet packed with the catalyst bed (maintained in the range 250-400 °C). The feed pulse vaporized in the front inlet upon injection, and was allowed to pass through the packed catalyst bed at space velocities in the range of 1-45 s^−1^ by helium as the diluent gas.^44^ The injection was immediately followed by separation using HP-PLOTQ column (Agilent, 19091P-QO4), and compounds in the effluent stream were identified using online Agilent 5975C Triple-Axis MS detector, and quantification was enabled by a quantitative carbon detector (QCD, Polyarc™) in conjunction with a flame ionization detector (FID).

## Acknowledgement

This work was funded by the generous support of the National Science Foundation through a CAREER Award (029504-003) to M.C.Y.C. and the Center for Sustainable Polymers, a National Science Foundation-supported center for Chemical Innovation (CHE-1901635). H.S. acknowledges support from the Camille and Henry Dreyfus Postdoctoral Program in Environmental Chemistry. Z.W. also acknowledge the generous support from the Research Foundation for the State University of New York (71272).

## Author contributions

Z.Q.W., H.S., N.H., D.S.P., P.J.D. and M.C.Y.C designed research; Z.Q.W., H.S., N.H., D.S.P., and Y.M. carried out experiments; Z.Q.W., H.S., N.H., D.S.P., G.K., P.J.D., and M.C.Y.C analyzed data; and Z.Q.W., H.S., G.K., P.J.D., and M.C.Y.C wrote the paper.

## Competing financial interests

The authors declare no competing financial interests.

## Data availability statement

All data generated and analysed during this study will be included in the published article (and its supporting information files).

## Biological materials availability statement

All plasmids and strains generated in this study are available by request.

## Notes

### Competing Interest Statement

The authors have declared no competing interest.

