## Supplemental information for "A Hybrid Cellular-Heterogeneous Catalyst Strategy for the Production of Olefins from Glucose"

| **Supplementary materials and methods** |  |
| --- | --- |
| *Commercial materials* | S2 |
| *Transitive clustering of thiolase sequences* | S2 |
| *Construction of plasmids* | S3 |
| *Protein expression and purification* | S4 |
| *Enzyme screening assays* | S5 |
| *Enzyme assays with purified proteins* | S6 |
| *Steady-state kinetic analysis of pathway enzymes* | S6 |
| *MsFabG4 and MaoC stereochemical analysis* | S7 |
| *Deletion of fadE from E. coli MG1655(DE3)* | S7 |
| *Extraction and analysis of fatty acids by GC-MS and LC-MS* | S7 |
| *Heterogeneous catalyst screening* | S8 |
| **Supplementary results** |  |
| *Table S1. Strains, plasmids, oligonucleotides, gBLOCKS, and Accession IDs* | S10 |
| *Figure S1. Genomic context of PaaJ-like thiolase clusters* | S19 |
| *Figure S2. In vitro screening of PaaJ1-9 thiolases* | S20 |
| *Figure S3. Steady-state kinetic characterization of PaaJ5, PaaJ7, and PaaJ9 with various acyl-CoAs* | S21 |
| *Figure S4. In vitro characterization of enzymes in the PaaJ pathway* | S22 |
| *Figure S5. Thioesterase screening and characterization* | S24 |
| *Figure S6. In vitro reconstitution of the PaaJ thiolase pathway* | S25 |
| *Figure S7. Characterization of the PaaJ-dependent fatty acid biosynthesis pathway in engineered E. coli.* | S26 |
| *Figure S8. Heterogeneous catalyst screening for 3-hydroxyoctanoic acid* | S27 |
| **Literature Cited** | S32 |

**Materials and Methods**

**Commercial materials.** Luria-Bertani (LB) Broth Miller, LB Agar Miller, Terrific Broth (TB), and glycerol were purchased from EMD Biosciences (Darmstadt, Germany). Carbenicillin (Cb), kanamycin (Km), spectinomycin (Sp), D-glucose, isopropyl-β-D-thiogalactopyranoside (IPTG), dithiothreitol (DTT), diethyl ether, methanol, 4-(2-hydroxyethyl)-1-piperazineethanesulfonic acid (HEPES), magnesium chloride hexahydrate, sodium chloride, sodium phosphate dibasic, Tween-20, hydrochloric acid, and sodium hydroxide were purchased from Fisher Scientific (Pittsburgh, Pennsylvania). Adenosine triphosphate sodium salt (ATP), nicotinamide adenine dinucleotide reduced form dipotassium salt (NADH), nicotinamide adenine dinucleotide 2′-phosphate reduced tetrasodium salt (NADPH), nicotinamide adenine dinucleotide hydrate (NAD^+^), acetyl-coA, butyryl-CoA, hexanoyl-CoA, octanoyl-CoA, decanoyl-CoA, dodecanoyl-CoA, malonyl-CoA, crononyl-coA, 3-oxobutyryl-CoA (acetoacetyl-CoA), 3-oxohexanoyl-CoA, 3-hydroxyl butyryl-CoA, 3-hydroxyloctanoic acid, 3-hydroxyldecanoic acid, 2-hydroxylhexanoic acid, trimethylsilyl-diazomethane (TMS-DAM), lysozyme, β-mercaptoethanol (βME), acetonitrile (LC/MS-grade) were purchased from Sigma-Aldrich (St. Louis, Missouri). Butyric acid, hexanoic acid, octanoic acid, decanoic acid, dodecanoic acid, tetradecanoic acid, pentadecanoic acid, hexadecenoic acid, formic acid and imidazole were purchased from Acros Organics (Morris Plains, New Jersey). Restriction enzymes, T4 DNA ligase, Antarctic phosphatase, Phusion DNA polymerase and Taq DNA ligase were purchased from New England Biolabs (Ipswich, MA). Deoxynucleotides (dNTPs), were purchased from Invitrogen (Carlsbad, California). **Oligonucleotides were purchased from Integrated DNA Technologies (Coralville, Iowa), resuspended at a stock concentration of 100** µM in water and stored at -20 °C. DNA purification kits and Ni-NTA agarose were purchased from Qiagen (Valencia, California). Complete EDTA-free protease inhibitor was purchased from Roche Applied Science (Penzberg, Germany). Acrylamide/bis-acrylamide (30%, 37.5:1), electrophoresis grade sodium dodecyl sulfate (SDS), N,N,N',N'-tetramethyl-ethane-1,2-diamine (TEMED), and ammonium persulfate were purchased from Bio-Rad Laboratories (Hercules, California). PageRuler Plus Pre-stained Protein Ladder was purchased from Fermentas (Glen Burnie, Maryland). Amicon Ultra 10,000 MWCO centrifugal concentrators, MultiScreen_HTS_ 0.22μm filter plates, and Milli-Q Gradient water purification system were purchased from Millipore (Billerica, Massachusetts). Parafilm M was purchased from Penchiney Plastic Packaging (Akron, Ohio). GC vials and glass inserts were purchased from National Scientific (Alexandria, Virginia). 0.2 µm syringe filter was purchased from Waters Corporation (Milford, Massachusetts).

**Transitive clustering of thiolase sequences.** Thiolase_N family protein sequences (14,120) were downloaded from the Pfam website in the FASTA format. CD-HIT program [*1*] was used to reduce the protein family members to representative sequences of each cluster with a cutoff value at 60% identity. About 500 representative sequences were generated by this method. All-versus-all BLAST (Basic Local Alignment Search Tool) program was used to align all the representative sequences pair-wise to compare the similarity between any of the two input sequences [2]. A similarity network of all the representative sequences was built using the Blast2Similarity Graph plugin in Cytoscape 2.8 [*3*]. The resulting similarity network identified a branch of putative PaaJ-like thiolases from various Actinobacteria. In order to obtain more sequences from this branch, all sequences in the clusters of putative synthetic thiolases were compared pair-wise by all-versus-all BLAST to identify PaaJ-like thiolases from species other than Actinobacteria. The resulting file was used to generate a second similarity network by the same method. PaaJ1-9 proteins from the proteobacteria cluster were selected for the subsequent cloning.

**Construction of plasmids.** Plasmids were all constructed by Gibson assembly [4] except where otherwise noted. All PCR amplifications were carried out with Phusion polymerase or Pt Taq HF using oligonucleotides listed in **Table S1**. Double strained DNA fragments (gBLOCKs) were synthesized by Integrated DNA Technologies (**Table S1**) and resuspended at a stock concentration of 20 ng/µL in double-distilled water. All primers were designed to contain 25 bp overlap with the neighboring DNA fragments. Vectors were digested with appropriate restriction enzymes then mixed with inserts, either gel-purified PCR products or synthetic gBLOCKs. 1.5x Gibson mix was added into the mixture, and the Gibson reaction was carried out at 50ºC for 1 h in a thermocycler. The resulting mixture was transformed immediately into the chemically competent *E. coli* DH5α by heat shock. Details of individual plasmid constructed is listed below:

*pET16b-MspaaJ1.* The *MspaaJ1* coding sequence was PCR amplified from the genome of *M. smegmatis* using the pET16b *MspaaJ1* f and r primer pair. The PCR product and the pET16b vector were digested with NdeI and XhoI. The cut vector and the PCR product were ligated using the T4 DNA ligase.

pET16b-*Msf*abG4. The *MsfabG4* coding sequence was PCR amplified from the genome of *M. smegmatis* using the pET16b *MsfabG4* f and r primer pair. The PCR product and the pET16b vector were digested with NdeI and PspXI. The cut vector and the PCR product were ligated using the T4 DNA ligase.

*pET24a*-paaJ1-9 and other pET24a-based constructs for heterologous protein expression were assembled by Gibson reaction with synthetic gBLOCK fragments and pET24a digested by NdeI and XhoI.

*pCDF2-paaJ9 operon.* The full-length *paaJ9* was PCR amplified from pET24a-paaJ9 using the pCDF2 *paaJ9* f and r primer pair. The full-length *fabG9* and *maoC9* were obtained similarly. The three genes were inserted simultaneously into the pCDF2 vector digested by BamHI and HindIII by Gibson assembly.

*pCDF2-paaJ9.MsfabG4.maoC9.* gBLOCK fragments of *MsfabG4* f1 and f2 were inserted into the SacI-KpnI sites of the pCDF2-*paaJ9* operon vector by Gibson assembly.

*pCDF2-paaJ9.MsfabG4.mfe2.* The plasmid *pCDF2-paaJ9.MsfabG4.maoC9* was digested with KpnI and HindIII to remove the *maoC9* gene. gBLOCKs of *mfp2* f1 and f2 were directly inserted into the cut vector by the Gibson assembly.

*pCDF2-paaJ9.MsfabG4.echR.* The plasmid *pCDF2-paaJ9.msFabG4.maoC9* was digested with KpnI and HindIII to remove the *maoC9* gene. gBLOCKs of *echR* f1 and f2 were directly inserted into the cut vector by the Gibson assembly.

*pCDF2-paaJ1-8. MsfabG4.maoC9.* gBLOCKs of *paaJ1-8* were first inserted into the NdeI/XhoI sites of the pET24a vector to construct pET24a-*paaJ1-8* expression vectors. These vectors were later used as PCR templates for amplifying *paaJ1-8* using the primer pair pCDF2 *paaJ1-8* f and r. The plasmid backbone was generated by digestion with BamHI and SacI.

*pCDF2-paaJ9. fadJ.* Plasmid pCDF2-*paaJ9.MsfabG4.maoC9* was digested with SacI and HindIII. The *fadJ* gene was PCR amplified from the *E. coli* genome. The digested vector and the *fadJ* were ligated by Gibson assembly.

*pCWori-Tdter.tesB.* Plasmid pCWori-*Tdter.adhE2* was digested with EcoRI and KpnI to remove the *adhE2* gene [*5*]. The *tesB* gene was PCR amplified from the genome of the DH10B strain using the primer pair pCWori *tesB* f and r. The *tesB* gene was inserted into the cut vector by Gibson assembly.

*pCWori-Tdter.TE constructs (CpfatB1, AtTe, UcfatB1, tesA(L109P)).* pCWori-*Tdter.tesB* was digested by EcoRI and KpnI to remove the *tesB* gene. The gBLOCKs for thioesterases were assembled by Gibson followed by PCR amplification with the corresponding primer pairs. The purified PCR product was directly inserted into the cut vector by Gibson assembly.

*pZW-AtTe.* pESC-Leu was digested with EcoR1 and KpnI, cutting within the leu marker but resulting in a functional prokaryotic promoter upstream (-10, TATAAT; -35, ATGATA) while retaining both origins and the Ap^r^ marker. The PCR product of *AtTe* amplified from pET24a-*At*TE was inserted into the digested vector by Gibson assembly.

*pZW-Tdter.AtTe.* The pZW-*AtTe* plasmid was digested by the EcoRI enzyme. The *Tdter* gene was PCR amplified from the pCWori-*Tdter*.*AtTe* plasmid as template using the primer pair pZW *Tdter* f and r. The cloning was carried out by Gibson assembly.

**Protein expression and purification.** Terrific Broth (50 mL) containing kanamycin (50 mg/L) in a 250 ml baffled shake flask was inoculated at OD_600_ = 0.05 with an overnight TB culture of freshly transformed *E. coli* BL21(DE3) containing the appropriate pET24a or pET16b plasmids. The cultures were grown at 37°C at 200 rpm to OD_600_ = 0.6 and were induced with IPTG (0.1 mM). For PaaJ1, 2, 3, 7 and 9, the temperature was reduced to 30°C. Cell pellets were harvested 3 h post-induction. For PaaJ4, 5, 6, and 8, proteins were expressed at 16ºC overnight post-induction. Cells were collected by centrifugation at 15,000 × *g* for 20 min at 4ºC and stored at -80ºC for further analysis.

Frozen cell pellets were thawed and resuspended in 5 ml Ni-NTA lysis buffer (50 mM sodium phosphate, dibasic, 300 mM sodium chloride, 10 mM imidazole, protease inhibitor mixture pH 8.0 at 4ºC) with 0.2 mg/ml lysozyme on ice for 30 min. The cells were homogenized by sonication using the Misonix sonicator 3000 (Farmingdale, New York, United States) with a microprobe for 2 min with 10-s pulse and 10-sec intervals. Total lysate was passed through a 0.2 µm syringe filter before and then incubated with 3 mL Ni-NTA resin at 4ºC for 1 h. The purification process was carried out either by gravity-flow or using a fast performance liquid chromatography (GE Life Sciences, Marlborough, Massachusetts, United States) at a flow rate of 2 mL/min with 60 ml imidazole gradient from 10 mM to 250 mM using the lysis buffer and the elution buffer (50 mM sodium phosphate, dibasic, 1 M sodium chloride, 1% (*v*/*v*) Tween-20, 250 mM imidazole, pH 8.0 at 4ºC). 12 x 5 mL fractions were collected and analyzed by SDS-PAGE. Fractions containing pure protein were pooled and the buffer was exchanged to 100 mM HEPES, 1 mM magnesium chloride, 1mM DTT, and 20% (*v*/*v*) glycerol at pH 7.0 using an Amicon centrifugal filter unit at the appropriate cutoff size. Initial screening of thiolases PaaJ1-PaaJ9 (**Fig. S2**) was carried out on protein purified by gravity-flow Ni-NTA affinity chromatography with an initial wash at 10 mM imidazole followed by elution at 250 mM imidazole. The protein concentration was determined by its extinction coefficient at 280 nm on a Beckman-Coulter DU-800 spectrometer (Brea, California, United States). The purified protein was flash-frozen in liquid nitrogen before storing at -80ºC.

**Enzyme screening assays.**

*Thiolase screening assay.* PaaJ1-9 were screened for their ability to utilize butyryl-CoA as a starter unit using an endpoint assay monitoring the free CoA release in the presence of Ellman’s reagent (5,5’-dithiobis-(2-nitrobenzoic acid); DTNB) at 412 nm [*6*]. Substrates butyryl-CoA and acetyl-CoA were added at 100 µM. The concentration of DTNB was 300 µM. The buffer contained 50 mM HEPES and 5mM magnesium chloride, pH 7.0. The reaction was initiated by adding 1 µM purified thiolase and allowed to proceed for 60 min at 30ºC before recording A_412 nm_. The absorbance data was normalized with the lowest absorbance recorded with PaaJ8.

*Ketoreductase selectivity screening assay.* The relative ability of Hbd, PhaB, FabG9, or *Ms*FabG to carry out the ketoreduction of 3-oxobutyryl-CoA (acetoacetyl-CoA) compared to 3-oxohexanoyl-CoA was tested in an endpoint assay monitoring the oxidation of NAD(P)H at 340 nm at 30ºC [*7*]. The assay mixture contained the purified recombinant ketoreductase (0.1 μM), 3-oxoacyl-CoA (100 μM), and NAD(P)H (100 μM) in 50 mM HEPES, and 5mM magnesium chloride, pH 7.0. The reaction was initiated by adding 3-oxoacyl-CoA and allowed to proceed for 60 min 30ºC.

*In vivo screening assay for thioesterases.* LB broth (50 mL) containing carbenicillin (50 mg/L) and spectinomycin (100 mg/L) in a 250 ml baffled shake flask was inoculated at OD_600_ = 0.05 with an overnight culture in Terrific Broth plus 1.5% (*w/v*) glucose of freshly transformed *E. coli* DH1(DE3) strain containing the upstream plasmid: pCDF2-*paaJ9.fabG9.maoC9* or pCDF2, and the downstream plasmid: pCWori-*Tdter.TE* or pCWori. The cultures were grown at 37°C at 200 rpm to OD_600_ = 0.6 and were induced with IPTG (0.1 mM). After lowering the temperature to 30ºC, cultures were allowed to grow for an additional 3 d before quantification of fatty acids by GCMS as previously described.

**Enzyme assays with purified proteins.** UV-visible data were collected using a DU800 spectrophometer (Beckman Coulter, Brea, California) using a 1 cm-pathlength quartz cuvette. Initial rate data were collected for 10 min after initiation.

*Acetyl-CoA: acyl-CoA thiolase.* The thiolase assay was carried out in the carbon-carbon bond formation direction at 30ºC by monitoring the free CoA release in the presence of DTNB at 412 nm [*6*]. Substrate hexanoyl-CoA and acetyl-CoA were added at 100 µM. The concentration of DTNB was 300 µM. The buffer contained 50 mM HEPES and 5mM magnesium chloride, pH 7.0. The reaction was initiated by adding 1 µM purified thiolase.

*Acetoacetyl-CoA reductase.* The reduction of acetoacetyl-CoA was assayed by monitoring the oxidation of NAD(P)H at 340 nm at 30ºC [*7*]. The assay mixture contained the purified recombinant FabG9 or *Ms*FabG (0.1 μM), acetoacetyl-CoA (100 μM), and NAD(P)H (100 μM) in 50 mM HEPES, and 5mM magnesium chloride, pH 7.0. The reaction was initiated by adding acetoacetyl-CoA.

*3-Hydroxybutyryl-CoA dehydratase.* The dehydratase activity was measured in the forward direction by monitoring the formation of the double bond at 263 nm at 30ºC. The assay mixture contained 3-hydroxybutyryl-CoA (100 μM) in 50 mM HEPES and 5mM magnesium chloride, pH 7.0. The reaction was initiated by adding the purified MaoC9 (0.01 μM).

*Crotonyl-CoA reductase*. The crotonyl-CoA reductase activity was assayed in the forward direction by monitoring the oxidation of NAD(P)H at 340 nm at 30ºC [*8*]. The assay mixture contained the purified recombinant *Td*Ter (0.01 μM), crotonyl-CoA (100 μM), and NAD(P)H (100 μM) in 50 mM HEPES and 5 mM magnesium chloride, pH 7.0. The reaction was initiated by adding crotonyl-CoA.

*Acyl-CoA thioesterase.* The octanoyl-CoA thioesterase activity was assayed in the forward direction at 30ºC by monitoring free CoA release in the presence DTNB at 412 nm. The reaction mixture contained octanoyl-CoA (100 μM), DTNB (300 μM) in 50 mM HEPES and 5 mM magnesium chloride, pH 7.0. The reaction was initiated by adding the purified *At*TE protein (1 μM).

**Steady-state kinetic analysis of pathway enzymes**. Kinetic assays were carried out in a 1 cm-pathlength quartz cuvette on a DU800 UV-vis spectrophotometer (Beckman Coulter, Brea, California). Kinetic parameters (*k*_cat_ and *K*_M_) were determined by fitting the data using the Origin 6.1 to the equation: *V*_o_ = *V*_max_ [S] / (*K*_M_ + [S]), where *V*_o_ is the initial rate and [S] is the substrate concentration.

*PaaJ5, PaaJ7, and PaaJ9*. Activities of thiolases were measured in the forward direction by monitoring the free CoA formation at 412 nm at 30ºC in the presence of DTNB. The assay mixture (250 μL) contained 40 nM purified enzyme, 300 μM DTNB, and 1mM acetyl-CoA in potassium buffered saline (PBS) containing 137 mM sodium chloride, 2.7 mM potassium chloride, 10 mM sodium phosphate dibasic, 1.8 mM potassium phosphate monobasic, pH 7.4. The reaction was initiated by adding acetyl-CoA, butyryl-CoA, hexanoyl-CoA, octanoyl-CoA, or decanoyl-CoA at concentrations of 0.015625, 0.03125, 0.0625, 0.125, 0.25, 0.5, 1 mM.

*MsFabG4*. The activity of *Ms*FabG4 was measured by monitoring the oxidation of NADH at 340 nm at 30ºC. The assay mixture (250 μL) contained 0.1 μM purified *Ms*FabG4, 200 μM NADH, and 3 mM acetoacetyl-CoA in 50 mM HEPES with 5 mM magnesium chloride, pH 7.0. The reaction was initiated by adding acetoacetyl-CoA at concentrations of 0.0625, 0.125, 0.25, 0.5, 1, 2, 4 mM or NADH at concentrations of 2.5, 5, 10, 20, 40, 80, 160 μM.

*MaoC9.* The activity of MaoC9 in the reverse direction was measured by monitoring the hydration of crotonyl-CoA double bond at 340 nm at 30ºC. The assay mixture (400 μL) contained crotonyl-CoA at concentrations of 0.0625, 0.125, 0.25, 0.4, 0.5, 0.6, 0.7, 0.8, 1 mM in 50 mM HEPES with 5 mM magnesium chloride, pH 7.0. The reaction was initiated by adding 25 nM recombinant MaoC9.

*AtTE.* The activity of *At*TE was measured in the forward direction by monitoring the free CoA formation at 412 nm at 30ºC in the presence of DTNB. The assay mixture (250 μL) contained 1 μM purified *At*TE, and 300 μM DTNB in 50 mM HEPES with 5 mM magnesium chloride, pH 7.0. The reaction was initiated by adding acetyl-CoA, octanoyl-CoA, decanoyl-CoA, or dodecanoyl-CoA at concentrations of 0.03125, 0.0625, 0.125, 0.25, 0.5, 1, 2 mM.

***Ms*FabG4 and MaoC9 stereochemical analysis.** The acetoacetyl-CoA reductase and the 3-hydroxylbutyryl-CoA dehydratase were coupled and assayed in the reverse direction by monitoring the formation of NADH at 340 nm at 30ºC. The reaction mixture contained purified PhaJ, Crt, or MaoC9 (5 U), *Ms*FabG4 (0.1 μM), NAD^+^(2 mM) and crotonyl-CoA (1 mM) in in 50 mM HEPES and 5 mM magnesium chloride, pH 7.0. The reaction was initiated by adding crotonyl-CoA.

**Deletion of *fadE* from *E. coli* MG1655(DE3) using the λ red recombination system.** Deletion of the genomic *fadE* gene was achieved using the λ red recombination system [*9*]. Briefly, Km^R^ gene and flanking FRT sites were PCR amplified from the pKD4 plasmid using the *fadE* p1/p2 primers. The purified PCR products were transformed by electroporation into MG1655(DE3) cell bearing the pKD46 plasmid, which expressed the λ recombinase in the presence of the arabinose at 30ºC. Transformants resistant to kanamycin were selected, and the pKD46 plasmid was cured by streaking to single colonies on a nonselective plate at 37ºC overnight. The *fadE* mutation was confirmed by genotyping using the *fadE* KO1/2 primers. The helper plasmid pCP20 (Ap^R^) expressing the FLP recombinase was transformed into the Δ*fadE::kan^R^* cells to eliminate the Km^R^ at the *fadE* locus. Transformants were selected on an LB plus carbenicillin plate at 30ºC overnight. Colonies resistant to ampicillin were grown on a nonselective plate at 37ºC overnight to cure the pCP20 plasmid. The final Δ*fadE* MG1655 (DE3) was tested for the inability to grow on both Km- and Ap-selective plates.

**Extraction and quantification of fatty acids and derivatives.** Samples (500 µL) were removed from cell culture followed by centrifugation at 20,817 × *g* for 1 min using an Eppendorf 5417R centrifuge. The cleared media was mixed with 6N HCl (100 µL) and extracted with diethyl ether (500 µL) containing 200 mg/L pentadecanoic acid as the internal standard. For methyl transesterification, the organic layer (160 µL) was transferred to a GC vial glass insert and mixed with 2 methanol : 1 TMS-DAM (*v*/*v*) (40 µL) for at least 15 min at room temperature [*10*]. The samples were then quantified using a Trace GC Ultra (Thermo Scientific) with an HP-4MS column (0.25 mm x 30 m, 0.25 µM film thickness, J & W Scientific) and a DSQII single-quadrupole mass spectrometer (Thermo Scientific) using single-ion monitoring (74 and 88 *m*/*z* for fatty acids, 43 and 74 *m*/*z* for 3-hydroxy fatty acids) concurrent with full scan mode (*m*/*z* 35-100). The oven program was as follows: 50 ºC held for 3 min, ramped to 320 ºC at 30 ºC min^-1^, held at 320 ºC for 1 min. Samples were quantified relative to a standard curve of 8, 16, 31, 63, 125, 500 mg L^-1^ mixtures of butyrate, hexanoic acid, octanoic acid, decanoic acid, dodecanoic acid, tetradecanoic acid, hexadecanoic acid. Standard curves were prepared freshly for each run and normalized for injection volume using the internal pentadecanoic acid standard.

3-hydroxy fatty acids were quantified using LC-MS. Cell culture samples (200 μL) were cleared of biomass via centrifugation at 20,817 × g for 2 min. Cleared media (20 μL) was then diluted into water containing 0.25 g/L 2-hydroxyhexanoic acid as the internal standard (180 μL). Samples were filtered through a 96-well MultiScreenHTS plate (MilliporeSigma) before injecting onto an Agilent 1290 HPLC equipped with an auto-sampler and an Agilent EclipsePlusC18 RRHD column (50 × 2.1 mm). Solvent A was 0.5% *v*/*v* formic acid and solvent B was acetonitrile. The gradient was as follows: 0−1 min, 3% solvent B; 1− 6 min, 3−95% solvent B; 6−6.2 min, 95% solvent B; 6.2−7 min, 95−3% solvent B; 7−7.5 min, 3% solvent B at a flow rate of 0.6 mL/min. 3-hydroxy acids were quantified by mass spectrometry using an Agilent 6460 triple quadrupole MS with ESI source, operating in negative ion mode. Between 2-6 min, the following ions were monitored: 131.1 *m*/*z* (2-hydroxyhexanoic acid, internal standard); 159.1 *m*/*z* (3-hydroxyoctanoic acid); 187.1 *m*/*z* (3-hydroxydecanoic acid). Samples were quantified relative to a standard curve of 7.8125, 15.625, 31.25, 62.5, 125, 250, and 500 mg L^-1^ hydroxy acids. Standard curves were prepared freshly for each run and normalized for injection volume using the internal 2-hydroxyhexanoic acid standard.

**Heterogeneous catalyst screening.** Heterogeneous catalysts were screened using 8-hydroxyoctanoic acid on a pulsed microreactor system integrated with a GC-MS as described in the main text as well as in a published method [*11*]. 3-Hydroxyoctanoic acid was dissolved in water at a concentration of 5 mg/g, and passed over commercial catalysts (Silica-alumina (Si/Al = 13, Sigma Aldrich), H-Y (CBV 760, Si/Al = 60, Zeolyst), H-BEA (CP811C-300, Si/Al = 300, Zeolyst), γ-Al_2_O_3_ (Strem Chemicals), Nb_2_O_5_ (Sigma Aldrich), and ZrO_2_ (Sigma Aldrich) as vaporized pulses at temperatures in the range 250-400 ºC and a space velocity of 1 s^-1^ or 10 s^-1^ (typical injection volumes were 1 μL and up to 10 μL). The definition of space velocity is given by Eq.1, while conversion and product selectivities were calculated using Eqs 2 and 3, respectively. These experiments enabled shortlisting of Nb_2_O_5_ as the most olefin selective catalyst, which was subsequently tested for the fatty acid extracts (consisting of 3-hydroxyoctanoic, and 3-hydroxydecanoic acids). Analogous experiments with the extract mixture were performed using the same pulsed microreactor on Nb_2_O_5_ in the temperature range 325-400 ºC at a fixed space velocity of 45 s^-1^ to investigate the production of a mixture of heptenes and nonenes from the mixture (Figure 5C). For this case, the fermentation extract was fed directly to the reactor.

$SV \left[ s^{-1} \right]=\frac{Volumetric flowrate of diluent gas (He)[\frac{m^{3}}{s}]}{Volume of the catalyst bed [m^{3}]}$	(Eq.1)

$X [\%]=\left( 1-\frac{reactant concentration in the effluent}{Sum of reactant and product concentrations in effluent} \right)*100$	(Eq.2)

$S_{i}[\% C basis]=\frac{Total carbon in product i}{Total carbon in all products}*100$	(Eq.3)

**Supplementary Results**

**Table S1. Strains, plasmids, oligonucleotides, and sequences.** (A) Strains and plasmids used in this study. (B) Oligonucleotide sequences. (C) gBLOCKS used in the construction of synthetic genes. (D) Gene accession IDs.

**A. Strains and plasmids**

| **Strain** | **Genotype** | **Source** |
| --- | --- | --- |
| DH1 | *end*A1 *rec*A1 *gyr*A96 *thi*-1 *gln*V44 *rel*A1 *hsd*R17(r_K_^-^ m_K_^+^) λ^-^ | ATCC |
| BW25113 | *F- DE(araD-araB)567 lacZ4787(del)::rrnB-3 LAM- rph-1 DE(rhaD-rhaB)56, hsdR514 fhuA* | Ref. 2 |
| BL21(DE3) | F*^–^ omp*T *gal dcm lon hsd*S_B_(r_B_^-^ m_B_^-^) λ(DE3 [*lac*I *lac*UV5-T7 *gene 1 ind*1 *sam*7 *nin*5]) | Novagen |
| DP10(DE3) | F*^–^* *end*A1 *rec*A1 *gal*E15 *gal*K16 *nup*G *rps*L Δ*lac*X74 Φ80d*lac*ZΔM15 *ara*D139 Δ(*ara,leu*)*7697* *mcr*A Δ(*mrr*-*hsd*RMS-*mcr*BC) λ(DE3 [*lac*I *lac*UV5-T7 *gene 1 ind*1 *sam*7 *nin*5]) | Ref. 2 |
| CW2513(DE3) | λ(DE3 [*lac*I *lac*UV5-T7 *gene 1 ind*1 *sam*7 *nin*5]) | Ref. 2 |
| W3110(DE3) | F*^–^* *rph*-1 *INV*(*rrn*D, *rrn*E) λ(DE3 [*lac*I *lac*UV5-T7 *gene 1 ind*1 *sam*7 *nin*5]) | Ref. 2 |
| MG1655(DE3) | F*^–^ ilv*G- *rfb*-50 *rph*-1 λ(DE3 [*lac*I *lac*UV5-T7 *gene 1 ind*1 *sam*7 *nin*5]) | Ref. 2 |
| *ΔfadE* MG1655(DE3) | F*^–^ ilv*G- *rfb*-50 *rph*-1 *fadE* λ(DE3 [*lac*I *lac*UV5-T7 *gene 1 ind*1 *sam*7 *nin*5]) | This study |
| DH10B(DE3) | F*^–^* *end*A1 *rec*A1 *gal*E15 *gal*K16 *nup*G *rps*L *fhuA* Δ*lac*X74 Φ80d*lac*ZΔM15 *ara*D139 Δ(*ara,leu*)*7697* *mcr*A Δ(*mrr*-*hsd*RMS-*mcr*BC) λ(DE3 [*lac*I *lac*UV5-T7 *gene 1 ind*1 *sam*7 *nin*5]) | Ref. 2 |
| DH5α(DE3) | F*^–^* *end*A1 *gln*V44 *thi*-1 *rec*A1 *rel*A1 *gyr*A96 *deo*R *nup*G Φ80d*lacZ*ΔM15 Δ(*lac*ZYA*-arg*F)U169, *hsd*R17(r_K_^-^ m_K_^+^), λ(DE3 [*lac*I *lac*UV5-T7 *gene 1 ind*1 *sam*7 *nin*5]) | Ref. 2 |
| DH1(DE3) | *end*A1 *rec*A1 *gyr*A96 *thi*-1 *gln*V44 *rel*A1 *hsd*R17(r_K_^-^ m_K_^+^) λ(DE3 [*lac*I *lac*UV5-T7 *gene 1 ind*1 *sam*7 *nin*5]) | Ref. 2 |
| **Plasmid** | **Description** | **Source** |
| pCDF2 | *(2×Tac), lacI^q^, Sp^r^, ColDF13* | Millipore |
| pCDF2-*paaJ9.MsFabG4.maoC9* | *paaJ9.MsfabG4.maoC9 (2×Tac), lacI^q^, Sp^r^, ColDF13* | This study |
| pCDF2-*paaJ9.MsFabG4.maoC9* | *paaJ9.MsfabG4.maoC9 (2×Tac), lacI^q^, Sp^r^, ColDF13* | This study |
| pCDF2-*paaJ9.MsfabG4.mfe2* | *paaJ9.MsfabG4.mfe2 (2×Tac), lacI^q^, Sp^r^, ColDF13* | This study |
| pCDF2-*paaJ9.MsfabG4.echR* | *paaJ9.MsfabG4.echR (2×Tac), lacI^q^, Sp^r^, ColDF13* | This study |
| pCDF2-*paaJ1.MsfabG4.maoC9* | *paaJ1.MsfabG4.maoC9 (2×Tac), lacI^q^, Sp^r^, ColDF13* | This study |
| pCDF2-*paaJ2.MsfabG4.maoC9* | *paaJ2.MsfabG4.maoC9 (2×Tac), lacI^q^, Sp^r^, ColDF13* | This study |
| pCDF2-*paaJ3.MsfabG4.maoC9* | *paaJ3.MsfabG4.maoC9 (2×Tac), lacI^q^, Sp^r^, ColDF13* | This study |
| pCDF2-*paaJ4.MsfabG4.maoC9* | *paaJ4.MsfabG4.maoC9 (2×Tac), lacI^q^, Sp^r^, ColDF13* | This study |
| pCDF2-*paaJ5.MsfabG4.maoC9* | *paaJ5.MsfabG4.maoC9 (2×Tac), lacI^q^, Sp^r^, ColDF13* | This study |
| pCDF2-*paaJ6.MsfabG4.maoC9* | *paaJ6.MsfabG4.maoC9 (2×Tac), lacI^q^, Sp^r^, ColDF13* | This study |
| pCDF2-*paaJ7.MsfabG4.maoC9* | *paaJ7.MsfabG4.maoC9 (2×Tac), lacI^q^, Sp^r^, ColDF13* | This study |
| pCDF2-*paaJ8.MsfabG4.maoC9* | *paaJ8.MsfabG4.maoC9 (2×Tac), lacI^q^, Sp^r^, ColDF13* | This study |
| pCDF2-*paaJ9.fadJ* | *paaJ9.fadJ (2×Tac), lacI^q^, Sp^r^, ColDF13* | This study |
| pET24a | *(T7), lacI^q^, Km^r^, ColE1* | Novagen |
| pET16b | *(T7), lacI^q^, Km^r^, ColE1* | Novagen |
| pET24a-*paaJ1-His_6_* | *paaJ1-his (T7), lacI^q^, Km^r^, ColE1* | This study |
| pET24a-*paaJ2-His_6_* | *paaJ2-his (T7), lacI^q^, Km^r^, ColE1* | This study |
| pET24a-*paaJ3-His_6_* | *paaJ3-his (T7), lacI^q^, Km^r^, ColE1* | This study |
| pET24a-*paaJ4-His_6_* | *paaJ4-his (T7), lacI^q^, Km^r^, ColE1* | This study |
| pET24a-*paaJ5-His_6_* | *paaJ5-his (T7), lacI^q^, Km^r^, ColE1* | This study |
| pET24a-*paaJ6-His_6_* | *paaJ6-his (T7), lacI^q^, Km^r^, ColE1* | This study |
| pET24a-*paaJ7-His_6_* | *paaJ7-his (T7), lacI^q^, Km^r^, ColE1* | This study |
| pET24a-*paaJ8-His_6_* | *paaJ8-his (T7), lacI^q^, Km^r^, ColE1* | This study |
| pET24a-*paaJ9-His_6_* | *paaJ9-his (T7), lacI^q^, Km^r^, ColE1* | This study |
| pET16b-*His_6_-MspaaJ1* | *his-MspaaJ1 (T7), lacI^q^, Km^r^, ColE1* | This study |
| pET24a-*fabG9-His_6_* | *fabG9-his (T7), lacI^q^, Km^r^, ColE1* | This study |
| pET16b-*His_6_-MsfabG4* | *his-MsfabG4 (T7), lacI^q^, Km^r^, ColE1* | This study |
| pET24a-*maoC9-His_6_* | *maoC9-his (T7), lacI^q^, Km^r^, ColE1* | This study |
| pET16b-*His_6_-TdTer* | *his-TdTer (T7), lacI^q^, Km^r^, ColE1* | Ref. 2 |
| pET24a-*AtTe-His_6_* | *AtTe-his (T7), lacI^q^, Km^r^, ColE1* | This study |
| pCWori-*Tdter.UcfatB1* | *Tdter.UcfatB1 (2×Tac), lacI^q^, Ap^r^, ColE1* | This study |
| pCWori-*Tdter.tesA(L109P)* | *Tdter. tesA(L109P) (2×Tac), lacI^q^, Ap^r^, ColE1* | This study |
| pCWori-*Tdter.tesB* | *Tdter.tesB (2×Tac), lacI^q^, Ap^r^, ColE1* | This study |
| pCWori-*Tdter.CpfatB1* | *Tdter.CpfatB1 (2×Tac), lacI^q^, Ap^r^, ColE1* | This study |
| pCWori-*Tdter.AtTe* | *Tdter.AtTe (2×Tac), lacI^q^, Ap^r^, ColE1* | This study |
| pESC-Leu | *Yeast (2μ, gal1p, gal10p, leu2), Ap^r^, pUC* | Agilent |
| pZW-*AtTe* | *AtTe, Ap^r^, pUC* | This study |
| pZW-*TdTer.AtTe* | *TdTer. AtTe, Ap^r^, pUC* | This study |

**B. Oligonucleotide Sequences**

| **Name** | **Sequence** |
| --- | --- |
| pCDF2 *paaJ9* f | gagcggataacaatttcacacaggaaacaggatccaaaatatttaacgttaaggaggttcattatgcgttcaactcgccggg |
| pCDF2 *paaJ9* r | tacctccttatatttatagtctttgagctcttagcgctcgacgataacggt |
| pCDF2 *fabG9* f | gagctcaaagactataaatataaggaggtaagacc atgccctttgggcgccct |
| pCDF2 *fabG9* r | tacctccttatttttgctaggtttggtacctcatgctcccattaaggcctgg |
| pCDF2 *maoC9* f | ggtaccaaacctagcaaaaataaggaggtaagatag atgtcccgtcaatggcacgac |
| pCDF2 *maoC9* r | ggcgagatcaccaaggtagtcggcaaataaaagcttttacaggctttcccagcggc |
| pCWori *tesB* f | agttgaacgtttcgatcgtatttaagaattcaataattttgtttaactttaagaaggagatatacatatgagtcaggcgctaaaaaatttact |
| pCWori *tesB* r | tgacagcttatcatcgataagcttggtaccttaattgtgattacgcatcaccccttc |
| pCWori *CpfatB1* f | ggcagaagttgaacgtttcgatcgtatttaagaattcggaaacgattcatcatttaaggaggtataaatgttgttgaccgctatcactact |
| pCWori *CpfatB1* r | atgtttgacagcttatcatcgataagcttggtaccttacgtttttccggtcgaaatagc |
| pCWori *UcfatB1* f | tcatgtttgacagcttatcatcgataagcttggtaccttacaggtcggtgtgtttgcga |
| pCWori *UcfatB1* r | tcatgtttgacagcttatcatcgataagcttggtaccttacaggtcggtgtgtttgcga |
| pCWori *tesA(L109P)* f | agttgaacgtttcgatcgtatttaagaattcggatcaacgtacaataaggaggtctcaaaatgctggaatggaagccgaaac |
| pCWori *tesA(L109P)* r | tcatgtttgacagcttatcatcgataagcttggtaccttagacacgcggctcagc |
| pCDF2 *fadJ* f | tgtcaccgttatcgtcgagcgctaagagctctgcccaaaatccgggcaccccaggaggtccaatatggaaatgacatcagcgtttac |
| pCDF2 *fadJ* r | gatcaccaaggtagtcggcaaataaaagcttttattgcaggtcagttgcagt |
| pCWori *AtTe* f | agttgaacgtttcgatcgtatttaagaattccacccactattcacatctataaggtagttctaatgaaattcaagaaaaagttcaaaatcgg |
| pCWori *AtTe* r | tcatgtttgacagcttatcatcgataagcttggtaccttaaacattcgtcttgatcttgccc |
| pZW *Tdter* f | attgatgttgaaccttcaatgtagggaattctccaagcaggaggaaaagaagaggagtcgtaaatgatcgtcaagccaatggt |
| pZW *Tdter* r | accttatagatgtgaatagtgggtggaattcttaaatacgatcgaaacgttcaacttc |
| pCDF2 *paaJ1* f | ggataacaatttcacacaggaaacaggatccaaacaaaaaagagaataaggaggacattaatgctgacggcccc |
| pCDF2 *paaJ1* r | aacctccttattgactttgctccccgagctctcagcgttccaccaccgc |
| pCDF2 *paaJ2* f | ggataacaatttcacacaggaaacaggatcctgaaaagaacaccagaaatttaagagaggtcactaatgccgcatccgctg |
| pCDF2 *paaJ2* r | aacctccttattgactttgctccccgagctctcaacgttccaaaatcgccaca |
| pCDF2 *paaJ3* f | ggataacaatttcacacaggaaacaggatccgctcgtagcgagctaagaagttaaggaggtcatatatgaaacatggtcgtcgtgt |
| pCDF2 *paaJ3* r | aacctccttattgactttgctccccgagctctcacttctcgataatagcggtaacg |
| pCDF2 *paaJ4* f | ggataacaatttcacacaggaaacaggatccttataaataagaaaacgcagaggaggaaaagcataatgtctaataagaataatcacccgaatag |
| pCDF2 *paaJ4* r | aacctccttattgactttgctccccgagctctcacttctccagaatgcacgtga |
| pCDF2 *paaJ5* f | ggataacaatttcacacaggaaacaggatccaaggagatatacatatgaacacgagc |
| pCDF2 *paaJ5* r | aacctccttattgactttgctccccgagctctcatttttcaataatcatggtcacaccc |
| pCDF2 *paaJ6* f | ggataacaatttcacacaggaaacaggatccgattcagaataaaaaaggtaagaggaggaacactatgtccaatcgtaaaaccatgc |
| pCDF2 *paaJ6* r | aacctccttattgactttgctccccgagctctcagcgctcgatgatggct |
| pCDF2 *paaJ7* f | aattgtgagcggataacaatttcac |
| pCDF2 *paaJ7* r | aacctccttattgactttgctccccgagctctcacttttcgacaatcgccacc |
| pCDF2 *paaJ8* f | ggataacaatttcacacaggaaacaggatccggacaaacgcgtccaacaacgtaaggaggtatataatggctcaagcgaaaaagac |
| pCDF2 *paaJ8* r | aacctccttattgactttgctccccgagctctcaacgctccagaatcgcg |
| pZW_*Tdter* f | attgatgttgaaccttcaatgtagggaattctccaagcaggaggaaaagaagaggagtcgtaaatgatcgtcaagccaatggt |
| pZW_*Tdter* r | accttatagatgtgaatagtgggtggaattcttaaatacgatcgaaacgttcaacttc |
| pCDF *MsfabG4* f | tgtcaccgttatcgtcgagcgctaagagctcggggagcaaagtcaataagga |
| pCDF *MsfabG4* r | ttacctccttatttttgctaggtttggtacctcaggcacctaacattgcc |
| pET16a *MspaaJ f* | cagcagcggccatatcgaaggtcgtcatatggccaatgaatcgagacg |
| pET16a *MspaaJ r* | ttcgggctttgttagcagccggatcctcgagttaggcctccaggatcgc |
| pET16a *MsFabG4 f* | cagcagcggccatatcgaaggtcgtcatatgccgggatcgttcc |
| pET16a *MsFabG4 r* | tttcgggctttgttagcagccggatcctcgagttacgcccccagcatc |
| *fadE* P1 | ccatatcatcacaagtggtcagacctcctacaagtaaggggcttttcgtttgtgtaggctggagctgcttc |
| *fadE* P2 | ttaaataattagcggataaagaaacggagcctttcggctccgttattcatcatatgaatatcctccttagttcctattcc |
| *fadE* KO1 | gtgtaccggataccgcca |
| *fadE* KO2 | tgacggggctgttctcg |

**C. gBLOCKS used for synthetic gene construction**

| **Name** | **Sequence** |
| --- | --- |
| *paaJ9* f1 | cgttcaactcgccgggtcgcaatcctgggtggtaatcgtatcccgtttgcgcgtagcaacggtgcgtatgcaaccgcatccaaccaggcgatgctgacggctgcactggaaggcctgatcgaacgttatcgtctgcacggcctgcgtctgggcgaggtggtcgcgggtgcggttctgaagcacagccgtgacatgaatctgacgcgcgaatgtgttctgggtagccgtctgagcccgcagaccccggcctacgacatccagcaagcctgcggcaccggcctggaagccgcgctgctggttgcgaacaaaatcgcgctgggtcagattgattgcggtatcgccggtggtgtggacagcaccagcgatgcgccgattgcggtgaatgagggcctgcgccacattctgctgcaagctaaccgtggtaagagcctgggtgagcgtttgaaaccgtttctgaagttgcgtccgcacc |
| *paaJ9* f2 | ggtgagcgtttgaaaccgtttctgaagttgcgtccgcaccacctgaaaccggaactgccgcgtaacagcgagccgcgcacgggtctgtctatgggtgagcactgcgagcgtatggcgcacacgtggcgtgttggccgcgcagagcaggacgagctggcactgctgagccatcagaagctggctgctgcttacgcggaaggttggcaagacgatctgctgaccccgtttttgtccctggctcgtgataacaatctgcgcccggacctgagcctggagcaactggcgcgtttgaagccggcctttgatcgctccggccagggcacgctgaccgcaggtaatagcaccccgctgaccgatggcgcatctctggttctgctgggtagcgaggcctgggcgcaacagcaaggtctgcaagtcctggcctatttggtggatggtga |
| *paaJ9* f3 | agcaaggtctgcaagtcctggcctatttggtggatggtgaaaccgcagcggttgacttcgtcacgggtcgcgagggtctgttgatggccccggtgtatgcggttccgcgcctgttggcacgtaatggtctgacgttgcaggactttgactactacgaaattcatgaggcgttcgccgcccaggttttgtgcaccttgaaggcctgggaggacgcggattactgccgtgagcgcctgggtctggaagcgccgttgggtgcgattgaccgcagcaaactgaacgtccgcggtagcagcctggccgcaggtcatccgttcgcagcaaccggcggtcgtattttggcgaacctggcgaaactgctggcaacggctggcaaaggtcgtggtctgattagcatttgtgcggcgggtggccagggtgtcaccgttatcgtcgagcgc |
| *paaJ1* f1 | ttttgtttaactttaagaaggagatatacatatgatgctgacggccccgagcctgcgtcgcgtggcaatcatcggtggcacccgcattccgttcgcgcgtgccaataccgcctacgcggaagcgagcaaccaagatatgctgaccgcagccctgcaaggtttggtcgatcgttaccgtctgggcggtgaacgcctgggtgaggtcgtggcgggcgcggttttgaagcatagccgtgacttcaatttgacccgtgaggctgtcctgagcaccaccctgggccacgatacgccggcctatgatattcaacaagcgtgtggcaccggcctggaggcagcattgctggttgcgtctaagatcgcgctgggccaaattgacgcgggcattgcgggcggtgttgacacgaccagcgatgcaccgattgccctgaacgaaaaattgcgcaagctgctgctgaaagccaatcgtggtcgtaccacggctgagaaggtcaa |
| *paaJ1* f2 | tcgtggtcgtaccacggctgagaaggtcaaaaccctggcaggtattcgtccgtctatgctgttgcgtccggcgctgccgcgtaacgctgagccgcgtacgggtttgtctatgggtgagcactgcgagcaaatggcgaagcgttggcgcattccgcgtgcagctcaagacgagttgacggtcgcaagccaccgtaacctggcggcagcgtatggccgtggcttctttaacgatctgattgtcccgttccgcggtctgaccaaagacggcaacctgcgcccggacatcgatccggcgaagctggcggccatgccgccggtcttcgaccgtaaaggcggcacgatgacggcggcaaacagcacgccgctgaccgacggcgcagcggtggtcctgctggcgagcgaagagtgggcgcaagcgcgtaatctgccggtcctggcatacctgcgcacgggtgaagcggcggcggttgattttgttgatggcgaggaaggcct |
| *paaJ1* f3 | ggttgattttgttgatggcgaggaaggcctgctgatggccccggcttatgccgtgccgcgtatgctgggtaagttgggcattacgctgcaagattttgacttctacgagattcacgaggcctttgcggcccaggttttgagcaccttggcagcgtgggagagcccggattatgcgcgtgatcgcctgggtctggacgcgccgccgggcgcgattgaccgtgcgcgtctgaacgtcaacggtagcagcctggcgacgggtcacccgtttgcagcaaccggcggtcgtattctggccacgctggcaaaaatgctggcggagaagggtgcgggtcgtggcctgattagcatctgtgcggcgggcggtcaaggtgtggtggcggtggtggaacgcctcgagcaccaccaccaccaccactgagatccggct |
| *paaJ2* f1 | ttttgtttaactttaagaaggagatatacatatgccgcatccgctgccgccggtgcgccgtgttgccatcatcggtggtaatcgtattccgtttgcccgttctaataccgcatacgctaccgcaagcaaccaggacatgctgacctttacgttgcaaggcctgattgaccgctataacctgcacggtgagcgtctgggtgaagttgcggcgggtgcagtgattaaacactctcgtgacttcaatctgacccgtgaatccgtcctgtccaccaccctggcgaaagaaaccccggcgtatgatgtgcagcaagcgtgcggtacgggtttggagacggctatcctggtggctaataaaattgcactgggtcaaattgatgttggtatcgccggtggtgtggacacgacgagcgacgcgccgattggtgttaacgaaaacctgcgtaagattctgttggaggcaaaccgtggtcgttctccgg |
| *paaJ2* f2 | tgttggaggcaaaccgtggtcgttctccgggtcaacgtgttggtgcgctggcgaagctgcgtccgggcatgtttttcaagccgttgctgccgcgcaacggcgagccgcgtacgggtctgagcatgggtgagcattgcgaactgatggcaaagcgctggaatattagccgcgaggcacaggacgtgctggcgtacgacagccaccgcaagctggcggaggcgtacgctcgtggctttgttaatgacctgatgacgccgtatcgtggcctggcacgcgataacaacctgcgcagcgatctgacgctggaaaaactggcgagcctgaaaccggcatttgatcgcgacgcgggtacgctgaccgcgggtaatagcaccccgctgacggacggtgcgagcgcggtcttgctggcgagcgaggattgggccgcccgtcgtggtctgccggtcctggcttatctgagctggagcgagacggcagccgttgactatt |
| *paaJ2* f3 | gctggagcgagacggcagccgttgactattttgacaaaaaggagggtctgctgatggccccggcctacgctgttccgcgtatgttggcacgcgccggcctgacgttgcaggacttcgacttctacgaaattcatgaagcctttgcggcgcaggtgctgtgcaccctggcagcttggcaggatgatgaatattgtcgtacccaactgggcctggatggcccgctgggtgcaattgaccgcgagaagctgaacgttaatggtggtagcctggcgacgggtcatccgtttgcggcaaccggcggtcgcattgttgcgggtttggctaagatgctggcacaattggacaaaccggctggcacggcacgtggtctgattagcatttgtgccgcgggtggccagggtgttgtggcgattttggaacgtctcgagcaccaccaccaccaccactgagatccggct |
| *paaJ3* f1 | ttttgtttaactttaagaaggagatatacatatgaaacatggtcgtcgtgtggcgatcatcggcggctctcgcatcccgttcgcacgcagcaataccgcatatattggtaagagcaaccaggacatgctgacggcagcctttcgtggtgtggttgataagttccaactgcacggcgaggaactgggtgatgtcgtggcgggtgcagttgtcaagcacagccgcgactttagcctgtgccgtgagtccgttctgggtagcggcctgtccccgatgaccccggcatttgacattcaacaggcgtgcggtacgggtctggaagccgcgatcgtcgtggcgaataagatcgccctgggccagatcgaggttggtatggcgggtggtgtggataccacgacggatgccccgattgcggtgagcgagggtctgcgtaagatcctgctggaagcaaaccgtgcacgtgatacgaaaggtaagctggcggctt |
| *paaJ3* f2 | cacgtgatacgaaaggtaagctggcggctttgctgaagattcgtccgaagcatctggctccgctgccgccggcgaacgctgaaccgcgcaccaaaatgagcatgggcgctcactgcgaagtgatggcccagacgtggaacattagccgcgaggaacaagacgagctggctgtcagcagccacctgaaggcggccaaagcgtacgaagaaggttttttcgatgatctggtcgtgccgtttaatggcctgacccgcgacaacaatctgcgtgcagatagcagcgtcgaacgtttgcaaggtctgaaaccggctttcgaccgtaaaagcggtaagggcaccatgaccgcgggcaacagcacgccgctgacggatggtgccagcgcggttctgctggcgagcgaagattgggcaaaagcccgcggtatccagccgctggcgtacctgacctttagcgagaccattggcgttgact |
| *paaJ3* f3 | cacgtgatacgaaaggtaagctggcggctttgctgaagattcgtccgaagcatctggctccgctgccgccggcgaacgctgaaccgcgcaccaaaatgagcatgggcgctcactgcgaagtgatggcccagacgtggaacattagccgcgaggaacaagacgagctggctgtcagcagccacctgaaggcggccaaagcgtacgaagaaggttttttcgatgatctggtcgtgccgtttaatggcctgacccgcgacaacaatctgcgtgcagatagcagcgtcgaacgtttgcaaggtctgaaaccggctttcgaccgtaaaagcggtaagggcaccatgaccgcgggcaacagcacgccgctgacggatggtgccagcgcggttctgctggcgagcgaagattgggcaaaagcccgcggtatccagccgctggcgtacctgacctttagcgagaccattggcgttgact |
| *paaJ4* f1 | ttttgtttaactttaagaaggagatatacatatgtctaataagaataatcacccgaatagcccggttatcgagacgacggtcgttaaagcatctgacgcaaagagcacggacacggcgacgggcaaggccgcaacggaggagagcaaaaaaccgaccgttttcgacaccgcggtcaaatctgatgccaccgacgccaagaagaccgatgcgaaagcggcagaaccgaaagcagcgtctaccaaaaaggaagcggcaaaaaatagcgatgtcgcaccggctaagaagaccaataccgcagctaaaaaggcaagcaccagcaaactgtcctccggccagaatcgtgtcgcgatcctgggcggtaaccgtatcccgttcgcccgtagcaacggctcttacgcggacgtgtctaacacggatatgctgaccgctgcgctggatggtttggttgaacgttacaacctgcaagacgaaa |
| *paaJ4* f2 | tggttgaacgttacaacctgcaagacgaaaaaattggcgaggttgtcgcgggcgcagtcatgaagctgagccgtgatattaatctgacgcgcgaggcgaccctgaatacggccctgaacccgcacaccccgacctacgacatcagccaagcgtgcggcacgggtttgcaagccaccttcgcaagcgcgaacaaaatcgcgctgggtatcatcgatagcgccattacgggcggcgttgatagcacctccgacgcgccgatcgcggtgggtgacggcctgcgcaaagttttgattaaactgggtgcggcgaaagataataagcaacgcctgaaagcgttgatgggtctgaacccgaaagacctgctggacacgccgaagaacggtgaaccgcgtaccggcctgagcatgggcgatcaccaagcgattacgacgctggaatggaacattagccgtgaggcacaggataagctggctttcg |
| *paaJ4* f3 | gccgtgaggcacaggataagctggctttcgactcccatcagaatctgacccgcgcgtatgacgaaggtttctttgatgacctgattacgccgtataaaggtctgacccgtgataacaacctgcgcccggatacgaccctggagaagctggcgaagttgaagccggtgttcggtaagaagaacgctaaaccgaccatgaccgcagcgaacagcaccccgctgacggatggcgcgagctgcattctgctgggcaccgacgagtgggctgcggcgcatggtctgaaaccgctggcatatatcgttcatcaagagaccgcggctgtggacttcatcggtaaaagcggtgacaccgagggtctgttgatggccccggcgtacgcagtgccgcgcatgttggagcgtgcgagcctgacgttgcaggattttgatttctacgagatccacgaggcgttcgcgagccaggttctgagcaccctggccgcgtgggaagatgaa |
| *paaJ4* f4 | ctgagcaccctggccgcgtgggaagatgaaaagttctgcgaagagcgtctgggtctggacgcgccgctgggttctattgatcgttccaaactgaacgttaatggcagcagcctggcggcgggtcacccgtttgccgcgacgggcgcacgtatcctggcgacggcggcgaaactgctggatcaaaaaggcagcggtcgcgcattgattagcatttgtgccgcgggcggtcaaggcgtcactgcattctggagaagctcgagcaccaccaccaccaccactgagatccggct |
| *paaJ5* f1 | ttttgtttaactttaagaaggagatatacatatgaacacgagcaaaaccgttcgcaaagttgcaatcgttggctacaaccgcatcccgttcgcgcgtgcaaacacggcctacgcaagcgttggcaaccaagagatgatgaccgcggcactgaacggtctggttgacaaatacaacttgcagggccaactgctgggtgaggtggcaggcggtgcggtcattaagcatacgtacgataataacctgatccgcgaatgtgtgatgaagacggctctggacccggctacgccggcatgtgatttgcagcaagcgtgtgacaccggcatcgaaagcgcagtgtatattgcgaacaaaatcgcattgggccagatcgagagcggtattgcaggtggtgttgatagcatcagcgacattccgatggctgttagcgagaagctgcgtaaagtgttgctggaggcgcgtcaggctaa |
| *paaJ5* f2 | taaagtgttgctggaggcgcgtcaggctaaaagcctgggtggcaaaatcaaggcgttcctgaagctgagcccgaaggacctgagcccgctggttccgaaaaatgaagagcagcagaccggcctgagcatgggtggccacacggaaatcaccgccaagtactacaaaattagccgtgaggaccaggacaattttgcgttgaaaagccatctgaacatggcgaaggcgtatgatgagggttttttcaatgacatgattaccccgttcaatggtctggagaaggataacaacttgcgtaaagatagcagcattgagaagctggccaaactgaaaccggcgtttgacaaggttaatggtacgctgaccgcaggcaacagcacgccgctgaccgacggtgcctcctgcatcctgctggcaagcgaggagtgggctaaggaacgcggtctgccgattctggcctacattaccttcgcggaggcagcggcgattgagtacgttaa |
| *paaJ5* f3 | cgcggaggcagcggcgattgagtacgttaaaaaccagcagaatctgctgctggccccgctgtttgcggcgagccgtatgctggaaaaagcagaactgaatttgcaagattttgattactacgagattcatgaggcgttcgctgcccaagttctggcaacgctgaaaatttgggagagcccggaactgagtgctgaactgggtctgaaaaagacgctgggcgcaatcgaccgcgaaaagctgaatgttaagggtagcagcctggccgctgcacacccgttcgcagcgacgggtggtcgcatcattggtgtgatggcgaagttgctgaatgagaaaggtagcggtcgtggcttcgttagcatttgcgcggcaggtggtcagggtgtgaccatgattattgaaaaactcgagcaccaccaccaccaccactgagatccggctgctaacaaagcccgaaaggaagctgagttggctgctg |
| *paaJ6* f1 | ttttgtttaactttaagaaggagatatacatatgtccaatcgtaaaaccatgcgtccggtggcaattctggcaggtagccgtacgccgtttaccaaatcctttaccaattatagccgtaccagcaaccgtgaactgatgacggctaccgttcgtgacctggtggataaaacccaattgcgcggcgcattgctgggcgacgtgagcctgggtgccgtgatgaagaacgcgagcgactggaatttggcgcgtgagaccgtgctgggtgcaggcctggacccgcacacgccgggctacgatgtgcaacgtgcgtgtggtacgggtctggagaccgtggcacagatcgctctgaagatcgcgagcggtcaaatcgaaagcggtatcggtggtggtacggataccaacagcgatattgcgggtgttctgccgcacgagtttagctggattatgatggaagcgcaaaaggagaaaaccctgggtggtcgtctgaa |
| *paaJ6* f2 | aaaggagaaaaccctgggtggtcgtctgaaaaagtttgcagaactgaaactgcaatacctgaagccgcgtttcccgaacgtgcaagaaccgcgcacgggtaaaagcatgggtgaccacacggagatgatggttaaagattggatgattacccgtgaggcacaagacgagttggcctatcacagccatcagaacgcggcaaaagcgtacgccgagggtttctacaaggacctggtgttcgatttcaaaggtctgaaacaggatatttttgtccgcccggacaccaccctggaaaaactggccaagctgaaaccggcatttgacttcaacggcacgggtacgctgaccgcgggtaactccaccccgctgaccgatggtgcgagcgcggtgttgctgggcagcgaagagttcgcaaaagaaaaaaatttgccggttctggcatatttcgttgacgcagactacgcggcggtggactttgtcaag |
| *paaJ6* f3 | gcagactacgcggcggtggactttgtcaagggtgagggcctgctgatggccccgacccgtgcggttgcaaatctgctgcgtcgcaataacctgaccttgcaagactttgatttctacgagattcatgaagcgttcgcgggtcaagttctgtgtaccctgaaagcgtgggaagacgaggagtactgccgtacccaactgggctggtctaaagctctgggcagcatcgatcgcagcaagatgaacgttaaaggtggcagcttggcgctgggtcacccgtttgcggcgacgggcggtcgtatcgtggcaagcctggcaaagatgctggcacaaaagggtagcggtcgcggcttgatctccatttgcacggcgggtggtatgggtgtcgcagccatcatcgagcgcctcgagcaccaccaccaccaccactgagatccggct |
| *paaJ7* f1 | ttttgtttaactttaagaaggagatatacatatgtcccaaaataccgtgcgccgtgtggccattctgggtggtaaccgcattccgtttgctcgtagcaatacggcatactttaaagcgagcaattccgacatgctgacggcagcactgaatggtctggttgagcgctttaacctgcaaggtaagcgtatcggtgaggtcgtggccggcgcagtcttgaagcatagccgcgattttaacatgacgcgcgaagtcgttctgagcacggatctggcaccggagaccccggcgtacgatattcaaattgcctgcggtacgggcctgcaagcggcgttcgtggtggcaaataagatcgcgctgggtcaaatcgacgtgggtattgcgggtggcgtggatacgacctctgacgccccgatcgcagttggtgatggcctgcgcaaagtcctgctggagctgaatgttgccaagacgggtaaagatcgcctgaaagcgctgacgaa |
| *paaJ7* f2 | gggtaaagatcgcctgaaagcgctgacgaaaatcgacttcaagaagctgctggatgcaccgagcaacggcgagccgcgtacgggtttgagcatgggcgaacaccaggcgattacggccctggaatggggcattacccgcgaggcacaggacgaattggcggcgagcagccaccagaagttggcagcggcatacgaacgtggcttctttgacgacctgatgaccccgttcttgggcctgaatcgcgacaataatctgcgcccggacagcaccgtcgagaagctggcgaaactgaagccggtctttggtaagggtgagaccgccaccatgaccgcgggcaatagcaccccgttgaccgacggtgcaagcgtcgttctgttggcctccgaggagtgggccaaggagaacggtcatgaagtgctggcgtacctgagcttctccgagacggcagccgttgacttcattggtaa |
| *paaJ7* f3 | cgagacggcagccgttgacttcattggtaaaaatggcccgaaagaaggtctgttgatggctccggcctacgctgtgccgcgtatgctgaaacgcgccaacttgaaactgcaagacttcgacttttatgagattcacgaagcatttgcctcccaagtgctgtccaccctgaaagcatgggaggacgagaagttctgtaaagaacgtctgggtctggacgccccgctgggttccattgaccgtagcaaactgaatgtcaacggtagcagcctgggtgcaggtcacccgttcgctgccaccggcggtcgcattttggcgaccgctgcaaagctgattaatgaaaaaggtagcggtcgtgcactgattagcatctgtacggcgggcggcgaaggcgtggtggcgattgtcgaaaagctcgagcaccaccaccaccaccactgagatccggct |
| *paaJ8* f1 | ttttgtttaactttaagaaggagatatacatatggctcaagcgaaaaagaccagcgccccggttaagggtgcaagccagggcatccgccgcgtcgccgtgatcggcggtaaccgtattccgtttgcccgttctaacaccgcgtatagcaagattagcaatcaggagctgctgaccagcgcactgcgcggcctggttgatcgtttcaatttggatggtatgaagatgggtgaagtcgtcgcaggtgcggttattaaacactctcgcgatttcaacctgacccgcgaaagcgtgttgagctgtggtctggccccggaaaccccggcgtacgacatccaacaagcgtgcggtacgggtctggaggcagcgatcctggtggctaacaaaatcgccctgggtcaaattgaatgcggtattgcgggtggcacggatacgacctccgacgctccgatcggtgtcggtgagggtctgcgtgaaa |
| *paaJ8* f2 | cgatcggtgtcggtgagggtctgcgtgaaattctgttggacctgaaccgcgctaaaacgaccaaagagcgtttgaaaattttgggccgcttccgtccgagccatctggtgccggagatcccggaaaacggcgagccgcgcacgggcatgagcatgggcgaccattgccaggtgaccgcgaaagagtggagcattgctcgtgaggaccaggaccgtttggcctgggaaagccatcagaaactggctaaggcatacgaagaaggcttcttcgacgacctgatgaccccgatggcgggcctggacaaagacaacatcctgcgcccggataccacgctggagaagctggcaaccctgaagccgtgcttcgaccgtgagaatggtacgatgaccgcagcgaatagcaccgcactgacggacggtgcgtccgcagttctgctggcaagcgaggagtgggcgaaagctcataatatggacgttaaggcctggctgaatt |
| *paaJ8* f3 | ataatatggacgttaaggcctggctgaatttttccgaggttgcagcggtggacttcgtggataagaaggaaggcttgctgatggcaccggcctacgcggtcccgcgtatgctggagcgtgcaggtctgacgttgcaagactttgatttttacgagatccacgaggcttttgcagcacaagttctgtccacgttgaaagcatgggaagatccgggcttctgtcgcgaacgtctgggcctggaaaagccgctgggcaccatcgatcgcgacaaattgaacgtcaaaggcagcagcctggcgacgggccatccgttcgccgcgaccggcggtcgtattgttgctaccctggcaaagctgctggagcagaaaggtagcggtcgtggtctgatctctatctgtgcagcaggtggccagggcgttaccgcgattctggagcgtctcgagcaccaccaccaccaccactgagatccggctgctaacaaagcccgaaaggaagctga |
| *fabG9* f1 | gagctcaataattttgtttaactttaagaaggagatatacatatgccctttgggcgccctcttatgagtgatcgttatttgggtttcgcgaatagcaatctgggccgtcgcttggtggatgctctgggtttaccgcgtccggcgcctcttgaacgttggcaggctggccgcctgcgtccggtggaaggcgccctggtgttaggtggaggccctctggccaaacaggttgaagccatcgcaccgcggctcaccgatgaagtgtactcttttaatgcagacaatctgcaagcagaagcatgggtcgccggtctgggtccaaagattaaagccgtcgtcttcgacgctagccatctgtcagattccgacgctctgaaacagttgcgtgagttttttcaaccccttctgcgttcactggcgccgtgcgcacatgttttggtgttgggtcgtgctccggaggggttggatgatccgttggcgtctgtggcgcaacgtgcgctgg |
| *fabG9* f2 | gcgtctgtggcgcaacgtgcgctggaaggctttagccgtagcctggcgaaggaactgcgtaatggtgcaacggctcagttgctgtacgttgcaccgggtgcggaggaccaactggagggcgcactgcgcttttttctgtccccgaaatctgcttttattagcggccaagttctgcgtctgcaaggttgtacgagccaggttgaagactggacccgtccgctgggtggccgtcgcgctttggtcacgggtgccgcccgcggcatcggcgcagctatcgcggaaacgctggcgcgtgatggtgccgatgtgctgttgctggatgtcccgcaggccagcaaggacctggacgcactggcagcgcgtctgggtggcaaagccttgccgctggacatctgcgcaagcgacgcagccacccaactgctggcggctctgccggatggtatcgacatcgtggttcataacgctggtattacccgcgacaaaaccctggttaacatg |
| *fabG9* f3 | ccgcgacaaaaccctggttaacatgacacctgaatactgggatgctgtcttggcagtaaatttgaaagcaccgcaagtcctgacacaggctctgtatgacaacggtgcactgggcgaaaacgcccgtattactcttctggcgtccgtaagtgggattgctggaaatcggggtcaggcaaattacgcagcgagcaaagccggtttgattggtctggcgcaagcttgggcgccgcgtttagcggaacgtggcggttctatcaatgccattgcgcctggctttattgaaactcacatgactgctgcgatgcccatgggccttcgcgaggccggtcgtcggctcagctcactgggtcagggcggtagtcctcaggatgtggcggaagcgattgcatggttatcccagccaggcagcggatcggtgaatgggcaggtcctgcgtgtttgtggccaggccttaatgggagcatgaggtaccaataattttgtttaactttaagaa |
| *mfe2* f1 | aggcaatgttaggtgcctgaggtaccaaacctagcaaaaataaggaggtaagatagatggattcgcgcaccaagggcaaaaccgttatggaagtgggaggtgatggtgtcgccgtgatcaccctgatcaacccgcccgtcaatagtttgtcattcgacgttctttacaacttaaaaagcaattatgaagaggcgctttcgcgtaacgacgtgaaggcgattgttatcactggtgctaagggtcgtttctccgggggatttgatatttctggttttggggagatgcagaaggggaatgttaaggagcctaaggcaggttatatttccatcgacatcattactgacctgttagaagctgcacgtaagccgagtgttgctgctattgatggtctggctttaggtggtggtctggagttagccatggcttgccatgctcgtatctcagccccggcagctcagttagggttgcccgagctgcaattaggcgtgatccccggctttggcggcacccaacgcttgccacgccttgttggacttacaaaagcacttgaaatgattttgaccagtaagccagttaaggcagaggagggacacagtttgggcttgattgatgcggtggtgccacccgcggaactggtaacgactgctcgccgttgggcattagatattgttggacgccgtaaaccatgggtgtctagcgtgtctaaaaccgacaaattaccgccgctgggagaggcccgcgaaatccttactttcgcaaaggcgcaaacccttaagcgtgcaccgaatatgaagcatccgcttatgtgcttggacgctatcgaggttggcatcgtcagcgggccgcgcgctggcttggaaaaagaagctgaagttgcgtcgcaagtggttaagttggatacgactaagggactgatccacgtgtttttcagtcagcgcggaacagccaaagtacctggtgttacagaccgtggcttagtcccccgtaagatcaaaaaagttgcaattattggtggaggccttatggggtcgggtatcgcgacggcgttgatcttgagtaactacccagtgattttgaaggaagtaaacgagaaatttctggaggctggaattggacgcgtgaaagcaaatttgcagtcgcgcgtgcgtaaaggatcga |
| *mfe2* f2 | aattggacgcgtgaaagcaaatttgcagtcgcgcgtgcgtaaaggatcgatgtctcaggaaaaattcgagaagactatgtcattactgaaagggtcgttggattatgaatcgtttcgtgatgtcgacatggttattgaagccgttattgagaacatttcacttaagcagcaaatttttgcagatttggaaaagtattgcccacagcactgcatcttagcttccaacactagtacgattgatttaaacaaaattggggaacgtactaagtcgcaggatcgtattgttggcgcgcattttttttctcctgcccacatcatgccattattagagatcgttcgtacaaaccacacgagcgcgcaagttatcgtggaccttcttgacgtcggaaaaaagattaaaaagactcccgtcgttgttgggaattgcacggggttcgctgtgaatcgtatgttcttcccgtatacccaagccgctatgttcttagtggagtgcggtgccgatccctatttgattgatcgtgccatctccaagtttgggatgcccatgggacctttccgtctgtgcgaccttgtcggattcggagttgctatcgcgactgcgacacagttcattgagaatttttccgaacgcacgtataagagtatgatcattcctttaatgcaggaggataagcgtgcaggtgaagctacccgcaaaggtttctatttgtacgatgataaacgcaaagcgaaacctgatcccgaattaaagaaatatatcgagaaggcgcgcagcatttcaggcgttaaactggaccccaagctggccaatctttcggaaaaagatattatcgagatgactttctttccagtggtcaacgaggcatgtcgcgtcttcgctgaggggattgctgtgaaggcagcagacttggacatcgcggggattatgggaatgggtttccccccataccgtggcggtatcatgttttgggccgacagcattggcagtaaatatatctatagtcgtctggacgagtggagtaaagcatatggcgagtttttcaagccttgcgccttccttgctgagcgtggttcgaaaggggtattgttgtccgcccccgttaaacaagccagctcccgtcttagcttttatttgccgactaccttggtgatc |
| *echR* f1 | aggcaatgttaggtgcctgaggtaccaaacctagcaaaaataaggaggtaagatagatgcagaatgaaattaaaaaggtgtgtgtgattggagcgggagttatgggatcggggattgctgcactgatcgcgaacagttcacatcaagttgtattacttgacattttggacaaggactcgaacgatcccaataagatcgtaaaaaatagcgtgaaaaacttgcacaaacaaaaactgtctcctcttagcttcccagataaagtcaattttatcacgatcgggaatttggaacacgatttagacctgattaaggaatgcaacctggttattgaggtcatcgtggaaaagcttgaaattaagcatcagttatacaacaagattatcccgtaccttaaggaggacgctattattgcctcaaacaccagcacgttgccactgaagaagcttaaggtcaatttgccaaacaatatcaagtcacgctttgttatcacgcatttctttaacccgccccgctacatggaattggtggagcttattattgatcatactattaaagatgaagtgatcgtgaaagtctccgtgttcttgaccaaaatgctgggcaaaacaattatcaaatgtaatgatacaccggggttcattgcgaatcgtgtcggctgctttcttttggagttagtcgtccacaaagctattgcccagaaccttaattttgtcacaattgatcaaatcttctcccgttgcttggggctgcccaatacaggcatctttgggctttacgacttgattggtcatgacgtcatgaaattgatttcaagctccctgattagtgctcttccgaagagtgatgattatcatcgtatctatacgaatacaaaggtattcgataaaatgatcgagcataacttgatcggacgtaagggagaggggggcttttaccgtctttcagtcagcaacggcaaaaaaattaaagaggttatcaacatttctgatctgagctatcaccccgtacagaaagttgatatctctttcaactcactgaatgaattactttcctcaaattctatttatggtaagtttttcagtgagattatcacagaattctatatttatttaactagtttggtgccttcagtaactaacaacatctacgacattgaca |
| *echR* f2 | tttaactagtttggtgccttcagtaactaacaacatctacgacattgacacgacgatgaagttgggctatagttggcattacggcccttttgaattgctgaccattgccgtaaaaaacggatggaacttaatcatcaagaatgccgatctgatgaatattcctcttcctaaataccttgctagtaaagagtatcagaagatcgacaaacaaaagtttaatagcaaaaaagacttccttcaggagtcaaagattgttttgtcgaatgatagcgcgaatttaatccattattgtgagaatcttgtatttgtgatcaccacaaagatgaactctttgaatcacaatgtgttttacttgcttcaagaagcggtttcgaaggcggagaactacgggaaaaatctttatatttatcctcaggggaataatttcagtgccggtgccgacttaaaacttatcctgagctacatccaagacgggaactttcataacttagaaaatcttcttaagttggggcaacaaactatgcgttatttgaagtacagctcggttcacatcatttcctgtgcgcgcggtgttgcactggggggaggttgcgaacttttattgaattccagttatattgtcgcaaaccaggaactgaatgctgggcttatcgagttaggagttggattgattccaggctggacaggagttacagagatgttcgcccgctcaaacggaaataaaacaaagctgattcgcaatattaaaaatatcatcgagcagaacaaaacgtcttctgcggactacttcaaggcggactatggtatcaagaacatgcaagttaacatgaataaacactatattttagacgacgcactgaagttgcgcatttctaagaaaatcgtgagtatcccgaacaagattacgttaccaaaaattaatattgtgtctgaaattgacacgagtaaatataacgaactgcaaaacaaggttttgaataaatttcagaatattatcgataaacataacgagattagtgaggcggagttgttgacctacgagcgcgaaatgtttcttgagcttgctaagacgccgcaaactatcgagaagttgcaggcaatcgttggcagcttttatttgccgactaccttggtgatc |
| *AtTe* | atgaaattcaagaaaaagttcaaaatcggtcgtatgcacgttgatccgttcaattacatctccatgcgttacctggttgcgctgatgaatgaggtcgcctttgaccaagcagagatcctggaaaaagatattgatatgaagaacctgcgttggattatctactcttgggacattcagatcgaaaataacatccgcctgggcgaagaaattgaaatcaccaccattccgacccacatggacaaattttacgcctatcgtgacttcattgtggagtctcgtggcaacattttggcgcgtgcgaaagccacgttcctgctgatggacatcacccgtctgcgtccgattaaaattccgcagaatctgagcttggcctacggtaaagagaatccgatctttgacatctacgatatggagattcgtaacgacctggctttcattcgtgatattcaactgcgtcgtgcagacttggacaataattttcacatcaacaacgcggtgtacttcgacctgattaaagaaacggtggacatctatgacaaagacatctcttacatcaagctgatctaccgtaatgaaattcgcgataagaaacaaattcaagcgttcgctcgtcgtgaagataaaagcatcgattttgcactgcgcggtgaagacggtcgtgattattgcttgggcaagatcaagacgaatgtttaa |
| *UcfatB1* f1 | atgctggaatggaagccgaaaccgaagctgccgcaactgctggacgaccactttggtctgcatggtttggtgttccgtcgcaccttcgccattcgtagctatgaagttggtccggaccgttctaccagcatcttggcagtgatgaatcacatgcaggaggccacgctgaaccacgcgaaaagcgttggcattctgggcgatggtttcggcacgaccttggagatgagcaaacgtgacttgatgtgggtcgtgcgtcgcacgcacgtggcagtggagcgttatccgacctggggcgacacggtcgaggtggaatgctggattggtgccagcggcaataacggtatgcgtcgcgacttcctggttcgtgattgtaagaccggcgagatcctgacgcgttgcaccagcctgtccgttctgatgaacacccgtacccgccgcctgagcacgatcccggatgaggttcgtggcgaaatcggcccggcgttcattg |
| *UcfatB1* f2 | ttcgtggcgaaatcggcccggcgttcattgacaacgtggccgttaaggacgacgagattaaaaaattgcagaaactgaacgactccacggccgattacattcaaggcggtctgaccccgcgctggaacgatctggatgttaaccagcacgttaacaacctgaaatatgtggcgtgggtttttgagaccgtgccggacagcatttttgagagccaccatatctctagctttaccctggagtaccgtcgtgagtgcacgcgcgatagcgtgctgcgttccctgaccacggttagcggcggcagctccgaggcgggcctggtctgtgaccacctgttgcagctggaaggtggtagcgaagtcctgcgtgcgcgtaccgaatggcgcccgaaactgacggattcctttcgtggtattagcgtcattccggctgagccgcgtgtctaa |
| *CpfatB1* f1 | atgctgctggccgccatcacgaccgcgttcttggcggcagagaagcaatggatgatgctggaccgtaagccgaagcgtctggacatgctggaagatccgttcggcctgggtcgtgttgtccaggatggtctggtctttcgccaaaatttttccatccgtagctatgagatcggtgccgatcgtaccgcgagcattgaaaccgttatgaatcacttgcaagagaccgcattgaatcacgtcaaaaccgccggtctgagcaatgacggtttcggtcgtaccccggaaatgtataagcgtgatctgatctgggtggttgcaaagatgcaggtgatggtcaatcgttacccgacgtggggtgatacggtcgaagtgaacacctgggtggcaaagagcggtaagaacggcatgcgccgtgactggctgatcagcgactgtaacacgggtgagattctgacccgtgcaagcagcgtgtgggttatgatgaaccagaaaacccgcaagctgagcaagatcccggacgaagtgcgtcgtgagatcgagccgcatttcgttgacagcgccccggtcatc |
| *CpfatB1* f2 | ccgcatttcgttgacagcgccccggtcatcgaggatgacgaccgcaaattgccgaagctggatgagaagagcgcagatagcatccgcaagggtttgacgccgcgttggaatgacctggacgtgaaccaacatgttaataatgcgaagtacattggctggattctggaaagcaccccgccggaggtgctggagacccaagaactgtgctctctgaccctggaatatcgtcgtgaatgtggccgcgaaagcgtcctggagagcctgacggcggttgacccgagcggtgaaggctacggtagccaatttcagcatctgctgcgtctggaggacggcggtgagattgtgaaaggtcgtaccgagtggcgtccgaaaaacgccggtattaacggcgtggttccgagcgaggaaagcagcccgggtgactatagctaa |
| *tesA (L109P)* | atggcggacacgttattgattctgggtgatagcctgagcgccgggtatcgaatgtctgccagcgcggcctggcctgccttgttgaatgataagtggcagagtaaaacgtcggtagttaatgccagcatcagcggcgacacctcgcaacaaggactggcgcgccttccggctctgctgaaacagcatcagccgcgttgggtgctggttgaactgggcggcaatgacggtttgcgtggttttcagccacagcaaaccgagcaaacgctgcgccagattttgcaggatgtcaaagccgccaacgctgaaccattgttaatgcaaatacgtccgcctgcaaactatggtcgccgttataatgaagcctttagcgccatttaccccaaactcgccaaagagtttgatgttccgctgctgcccttttttatggaagaggtctacctcaagccacaatggatgcaggatgacggtattcatcccaaccgcgacgcccagccgtttattgccgactggatggcgaagcagttgcagcctttagtaaatcatgactcataa |
| *MsfabG4* f1 | tgtcaccgttatcgtcgagcgctaagagctcggggagcaaagtcaataaggaggttaataatgcctggttcgtttctcgccaaacagctgggcgttccccagccggaaacactgcgtcgttatcgtccgggcgatcctccattggcgggcagtttgttaattggtggtagcggtcgcgtggccgaacccctccggacggcgctggcagatgattacaatctggtctccaacaacattggtggtcgttgggctgattcattcggcggtgttgtctttgacgcaacagggattactgaagccgaaggtctgaaggaattatatacattctttacgccgctgttgcgtaacctggcgccttgtgcacgcgtagttgtcgtgggcaccacgccggccgaagccgggagcgttcatgcacaagtagtgcagcgtgccctggaaggcttcacgcgtagcctgggtaaaggtgcagcaccctggcgcgaccgcttaccgggtctgagcctggggcgccgtcaggctggtcggcatggatcacgtgtggatcacgcggtgcatccggtaggtcaggtgggatatgttgatgggcaggtttttcgtgttggggcagccgactcgacgcctcccgcagattgggataagccgttagatggaaaggtggctgtggtcaccggcgcggcacgtggcatcggagcgaccattgctgaagtatttgc |
| *MsfabG4* f2 | gagcgaccattgctgaagtatttgctcgcgatggcgcaacggtggttgccattgacgttgatggtgccgccgaagacttaaaacgtgtagcagataaagtgggtggcacagcgctgacgctggatgttactgcggatgatgcggtcgataagatcacggcgcacgttactgagcatcatggtggtaaagttgatatcctcgttaataacgctggtatcacccgcgataaactgctcgcgaatatggacgaaaaacggtgggatgcggtgattgccgttaatctgttagccccgcagcgccttacggaaggcctcgtgggtaacggtacaattggagaaggtggtcgcgtgatcggtctctcctcaatggctggcatcgccgggaaccgtggacagaccaattatgcaactactaaagcggggatgattggcctggcagaagccttggcaccagtactggctgataaaggtatcaccattaacgcagttgcaccgggctttatcgagaccaaaatgacagaagccatccctctggcgacgcgcgaggtcggtcgccgcctgaatagcctgttccagggtggtcagcctgtggatgtggctgaactcattgcatatttcgcaagtccggcttcgaacgcagttaccggaaacaccattcgcgtctgtggtcaggcaatgttaggtgcctgaggtaccaaacctagcaaaaataaggaggtaa |

**D. Gene accession IDs**

| **Protein** | **Source** | **Gene accession no.** |
| --- | --- | --- |
| *Ms*PaaJ1 | *Mycobacterium smegmatis* | WP_011726890 |
| *Ms*FabG4 | *Mycobacterium smegmatis* | WP_011726889 |
| PaaJ1 | *Acidiphilium cryptum* | WP_011942106 |
| PaaJ2 | *Burkholderia graminis* | ZP_02886425 |
| PaaJ3 | *Limnobacteria* sp. | ZP_01916313 |
| PaaJ4 | *Psychrobacter arcticus* | WP_011280582 |
| PaaJ5 | *Flavobacterium johnsoniae* | WP_012025306 |
| PaaJ6 | *Bdellovibrio bacteriovorus* | WP_011164517 |
| PaaJ7 | *Acinetobacter baumannii* | WP_000047940 |
| PaaJ8 | *Marinobacter aquaeolei* | WP_011785490 |
| PaaJ9 | *Pseudomonas putida* | WP_010951855 |
| PaaJ | *Mycobacterium flavescens* | WP_011891412 |
| PaaJ | *Mycobacterium sp. JLS* | WP_011557668 |
| FabG9 | *Pseudomonas putida* | AAN66208 |
| MaoC9 | *Pseudomonas putida* | AAN66207 |
| Mfe2 | *Arabidopsis thaliana* | NP_187342 |
| EchR | *Rickettsia prowazekii* | WP_015508398 |
| FadJ | *Escherichia coli* | WP_000426176 |
| *Td*Ter | *Treponema denticola* | WP_002681770 |
| *Cp*FatB1 | *Cuphea palustris* | AAC49179 |
| *Uc*FatB1 | *Umbellularia californica* | Q41635 |
| *At*TE | *Anaerococcus tetradium* | WP_004837416 |
| TesA(L109P) | *Escherichia coli* | WP_023567904 |
| TesB | *Escherichia coli* | WP_141066364 |

**Figure S1.** Genomic context of PaaJ-like thiolase clusters. (A) Three representative PaaJ clusters from *Mycobacterium* sp. (B) PaaJ1-9 clusters from Proteobacteria. *fabG*, 3-ketoacyl-CoA reductase; *maoC*, enoylacyl-CoA hydratase, *caiA* and *fadE*, enoylacyl-CoA reductase; *acrR*, putative transcription regulator; *caiC*, acyl-CoA ligase.

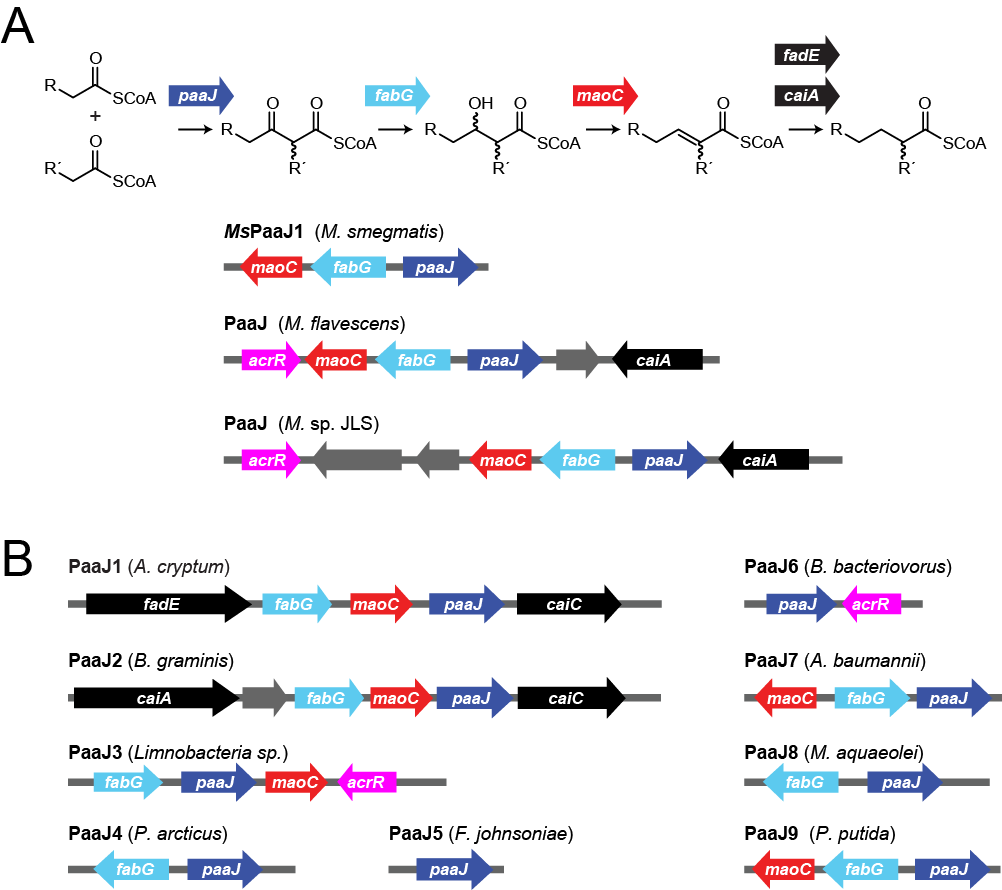

**Figure S2.** *In vitro* screening of PaaJ1-9 thiolases. (A) SDS-PAGE examining PaaJ1-PaaJ9 recombinant protein solubility in *E.coli* cell lysates. Expected MWs: PaaJ1, 45.1 kDa; PaaJ2, 46.4 kDa; PaaJ3, 45.2 kDa; PaaJ4, 55.6 kDa; PaaJ5, 45.9 kDa; PaaJ6, 46.4 kDa, PaaJ7, 45.7 kDa; PaaJ8, 47.1 kDa; PaaJ9, 45.5 kDa (U, uninduced cell lysate; S, soluble faction; I, insoluble fraction). (B) PaaJ1-PaaJ9 recombinant proteins used for the initial screening purified by gravity-flow chromatography. (C) *In vitro* screening of PaaJ1-9 activities. Butyryl-CoA (100 μM) and acetyl-CoA (100 μM) were used as substrates for PaaJ1-9 (1 μM) for monitoring condensation by free CoA release at 412 nm using DTNB and normalized for protein concentration.

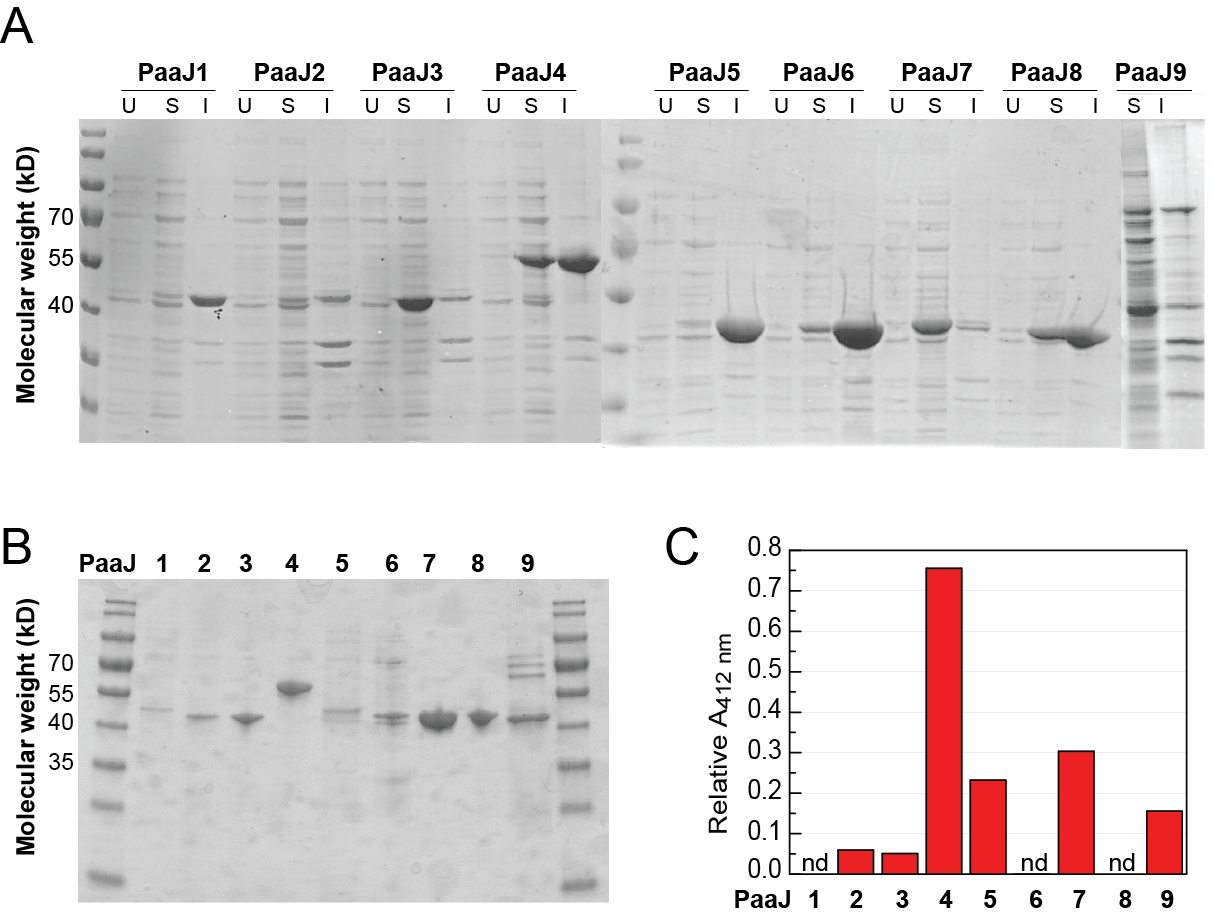

**Figure S3.** Steady-state kinetic characterization of PaaJ5, PaaJ7, and PaaJ9 with various acyl-CoAs. His_6_-tagged proteins used in these assays were purified by FPLC. Data are mean ± s,d, of technical replicates (n = 3). Tables contain *k*_cat_, *K*_M_, and *k*_cat_/*K*_M_ calculated by non-linear curve fitting to the Michaelis-Menten equation. Data are mean ± s.e. Error in *k*_cat_/*K*_M_ is obtained by propagation from the individual kinetic terms.

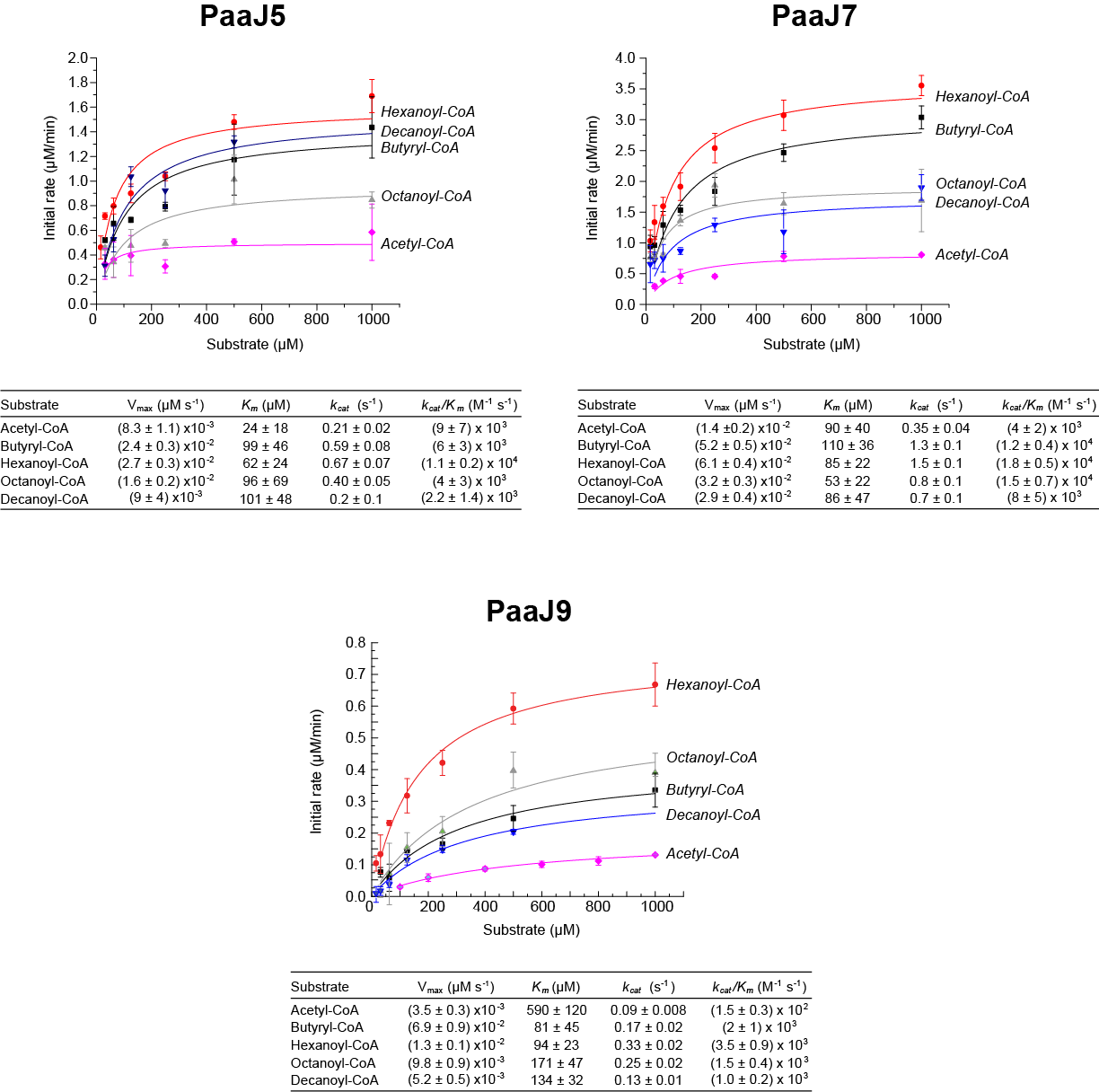

**Figure S4.** *In vitro* characterization of enzymes in the PaaJ pathway. (A) Purification and characterization of His_6_-tagged proteins (PaaJ9, 45.5 kD; FabG9, 47.8 kD; MaoC, 31.4 kD; *Td*Ter, 43.8 kD; *At*TE, 27.6 kD; *Ms*FabG4, 45.3 kDa) by FPLC. Data are mean ± s.d. of technical replicates (n =3). (B) Analysis of chain-length selectivity of *Ms*FabG4. When assayed on the C_4_ (3-oxobutyryl-CoA) compared to the C_6_ (3-oxohexanoyl-CoA) substrates, *Ms*FabG4 appears to prefer the C_6_ substrate. In comparison, canonical C_4_-selective ketoreductases, PhaB and Hbd, appear to prefer the C_4_ substrate. (C) Stereochemical analysis of the 3-hydroxyacyl-CoA intermediate. The dehydratases, PhaJ and Crt, selectively produce *(3R)*- and *(3S)*-hydroxybutyryl-CoA, respectively, when run in the reverse direction. Only coupling of *Ms*FabG4 to PhaJ results in NAD^+^ formation, suggesting that *Ms*FabG4 forms the (3*R*)-isomer. Likewise, coupling of the MaoC9 dehydratase to *Ms*FabG4 and not (3*S*)- producing Hbd results in NAD^+^ formation, which indicates that it prefers *(3R)*-hydroxybutyryl-CoA. Data are mean ± s.d. (n =3). (D) Steady-state Michaelis-Menten kinetic parameters for *Ms*FabG4 and MaoC9. Alignment of *Ms*FabG4 sequence with *E. coli* FabG also shows that D240 is found in place A36, which further suggests NADH selectivity because of the electrostatic repulsion with the phosphate group of NADPH. Kinetic data are mean ± s.d. of technical replicates (n =3). Table contains *k*_cat_, *K*_M_, and *k*_cat_/*K*_M_ calculated by non-linear curve fitting to the Michaelis-Menten equation. Data are mean ± s.e. Error in *k*_cat_/*K*_M_ is obtained by propagation from the individual kinetic terms. (SA, specific activity; nd, not detected)

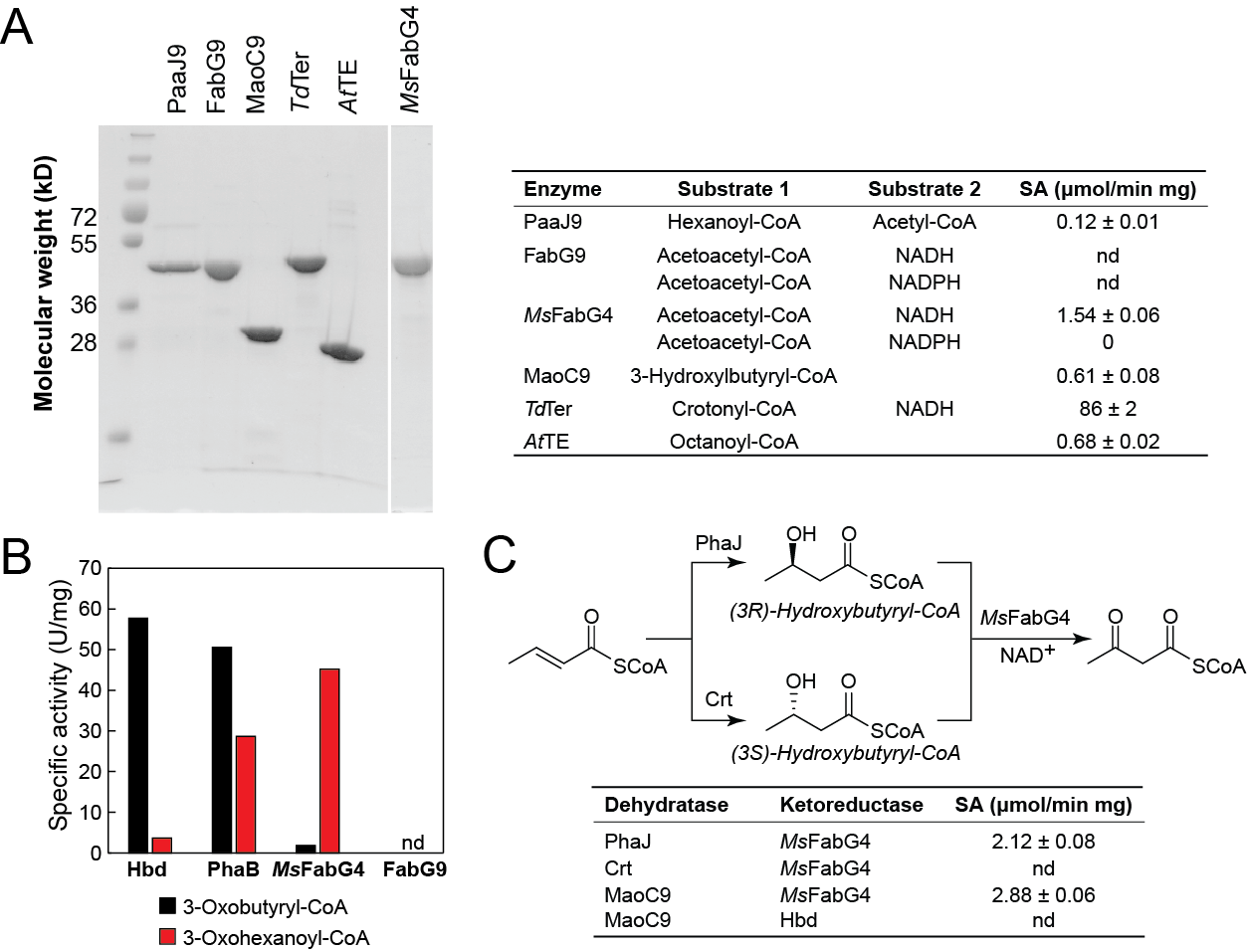

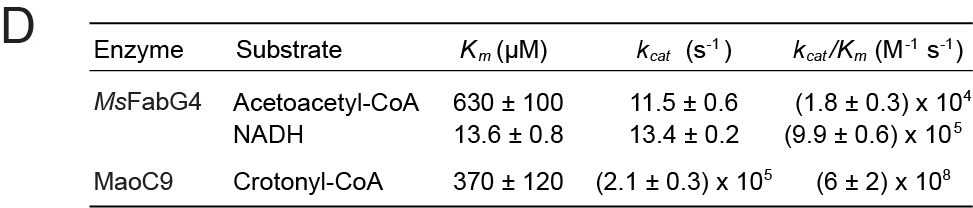

**Figure S5.** Thioesterase screening and characterization. (A) *E. coli* DH1(DE3) strains containing either no thioesterase (white, pCDF2 pCWori), thioesterase (black, pCDF2 pCWori-*Tdter.TE*), or upstream and downstream pathways (red, pCDF-*paaJ9.fabG9.maoC9* pCWori-*Tdter.TE*) were screened. After cell culture for 3 d, cell supernatants were acidified, extracted, and derivatized for GC-MS analysis. Fatty acids arising from strains with no upstream pathway (black) are likely derived from hydrolysis of acyl-ACPs. Fatty acids arising from strains with upstream and downstream pathways (red) may be derived also from hydrolysis of acyl-CoAs produced by PaaJ9-FabG9-MaoC9. For thioesterases where C4 data is omitted, no product was detected. Data are mean ± s.d. of biological replicates (n = 3). (B) Steady-state kinetic characterization of *At*TE. Initial screening of *At*TE on substrates of different chain length indicates that little activity is detected on butyryl-, hexanoyl-, and dodecanoyl-CoA (*left*). Michaelis-Menten parameters were then measured for octanoyl-, decanoyl-, and dodecanoyl-CoA (*right*). Data are mean ± s.d. of techical replicates (n =3). Table contains *k*_cat_, *K*_M_, and *k*_cat_/*K*_M_ calculated by non-linear curve fitting to the Michaelis-Menten equation. Data are mean ± s.e. Error in *k*_cat_/*K*_M_ is obtained by propagation from the individual kinetic terms.

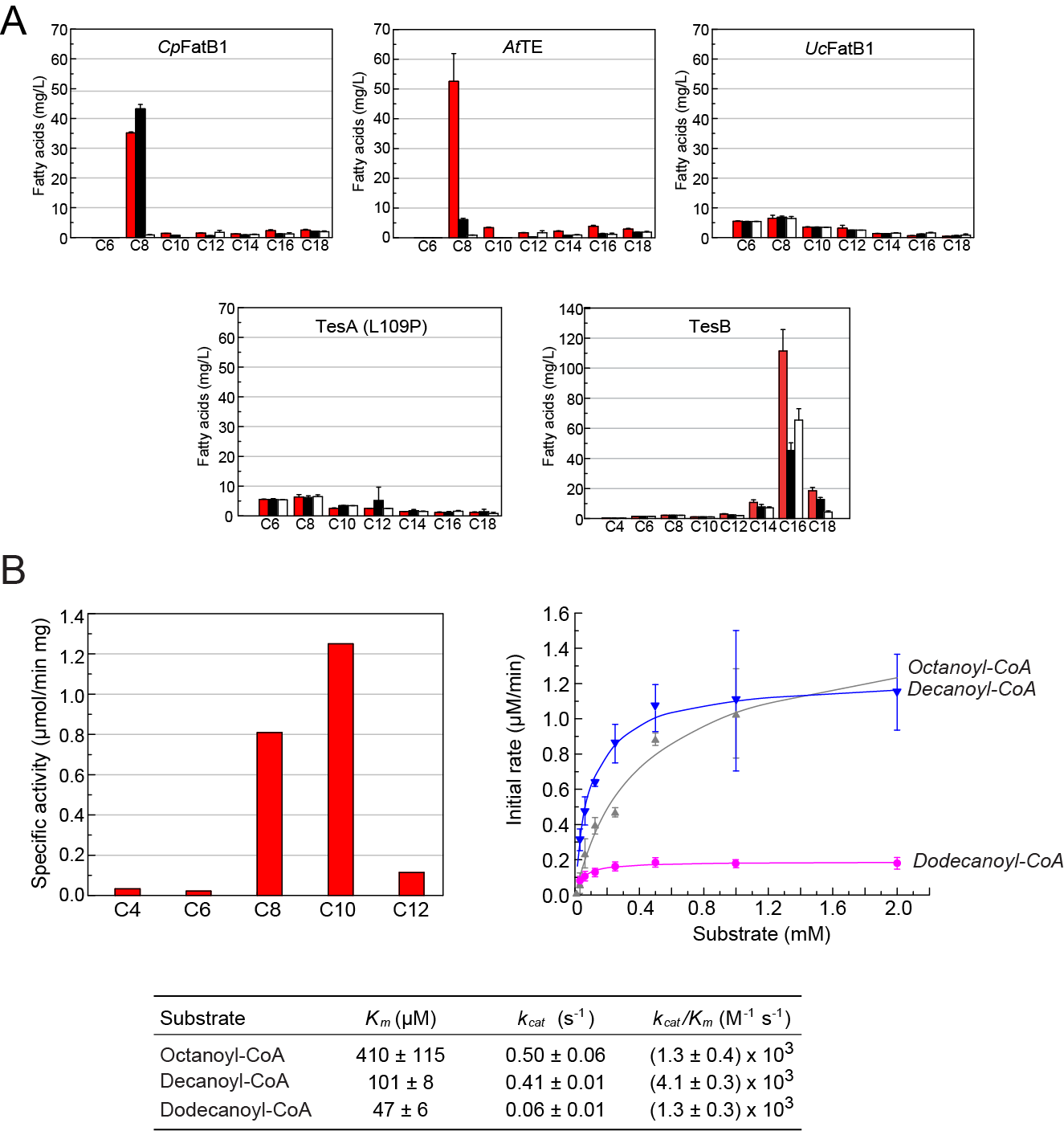

**Figure S6.** *In vitro* reconstitution of the PaaJ thiolase pathway. (A) *In vitro* single-step chain elongation from hexanoyl-CoA with acetyl-CoA and NADH using the five-enzyme system consisting of purified PaaJ9, *Ms*FabG4, MaoC9, *Td*Ter, and *At*TE. The reaction was analyzed by GC-MS after extraction and functionalization of the fatty acid products compared with a negative control omitting PaaJ9. The chromatogram shown is representative of three biological replicates (C8, octanoic acid; 3-OH C8, 3-hydroxy octanoic acid; C10, decanoic acid; 3-OH C10, 3-hydroxy decanoic acid; ISTD, pentadecanoic acid internal standard). (B) PaaJ9 titration of the iterative chain elongation system with purified PaaJ9, *Ms*FabG4, MaoC9, *Td*Ter enzymes, acetyl-CoA, and NADH at pH 7.0. The reaction was incubated at 37ºC for 1 h before acidification and extraction with diethyl ether. Samples were derivatized with TMS-DAM and quantified by GC-MS. Data are mean ± s.d. of technical replicates (n = 3). (C8, octanoic acid; C10, decanoic acid; C12, dodecanoic acid; C14, tetradecanoic acid, C16: hexadecanoic acid). (C) I*n vitro* iterative chain elongation reactions with PaaJ9 compared to PaaJ7 as described in B. Data are mean ± s.d. of technical replicates (n = 3).

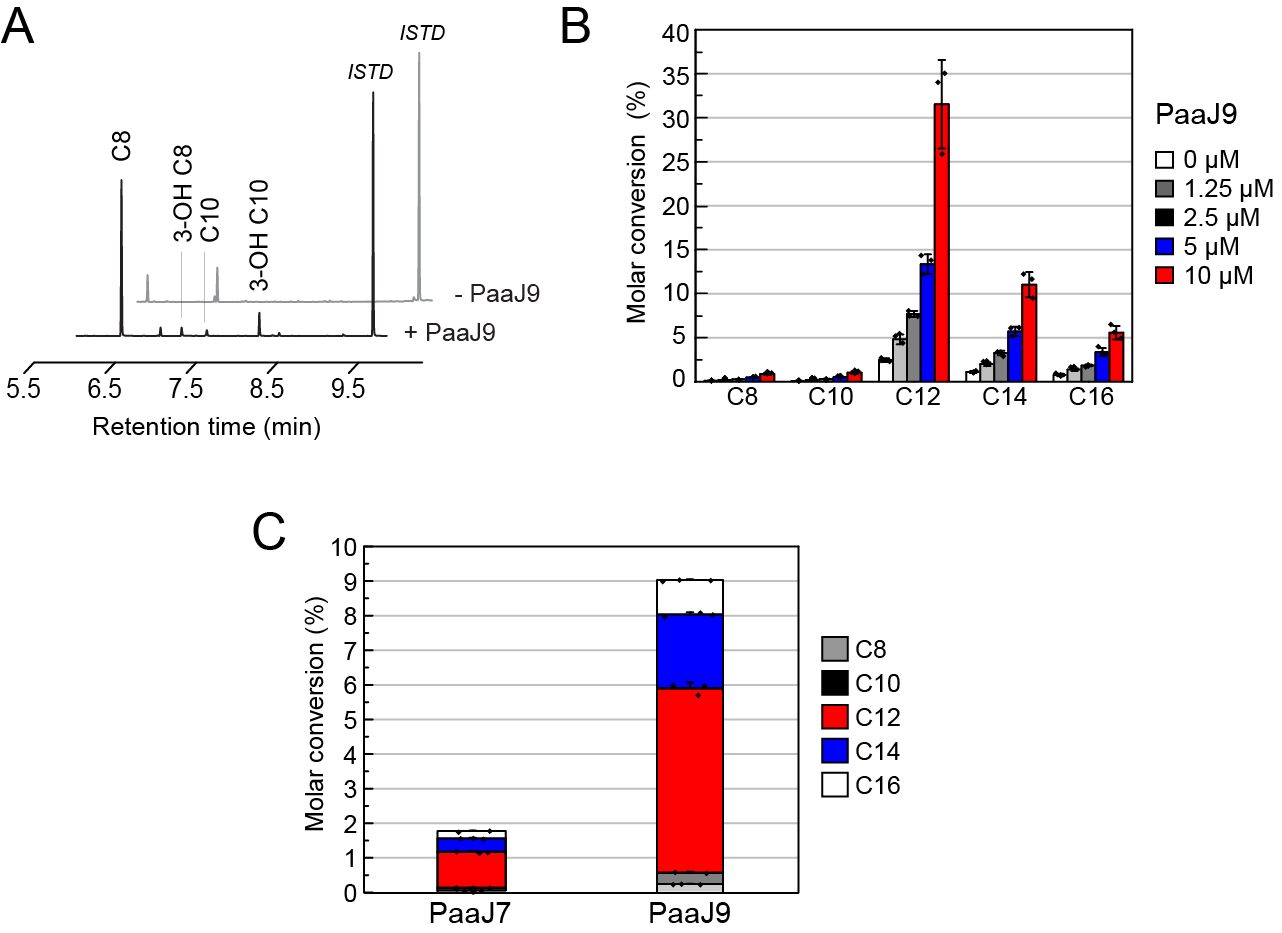

**Figure S7.** Characterization of the PaaJ-dependent fatty acid biosynthesis pathway in engineered *E. coli*. (A) Individual activity of pathway enzymes in cell lysates of strain **9** (*E. coli* MG1655 (DE3) *ΔfadE* pCDF2-*paaJ9.MsfabG4.maoC9* pZW-*Tdter.AtTe*) compared to the empty vector control. Data are mean ± s.d. of biological replicates (n = 3) (B) Growth curves for strains cultured microaerobically in shake flasks show a slight growth defect when *At*TE is expressed in the absence of the full PaaJ pathway. Strain **12** (*E. coli* MG1655(DE3) *ΔfadE* pCDF2-*paaJ9.fadJ* pZW-*Tdter.AtTe)* bearing the full pathway compared to the downstream pathway alone (*E. coli* MG1655(DE3) *ΔfadE* pCDF2 pZW-*Tdter*.*AtTe*) and an empty vector control. Data are mean ± s.d. of biological replicates (n = 3). (C) Fatty acid production in strain **12** compared to a control with an empty vector (pCDF2) in place of the upstream plasmid (pCDF2-*paaJ9.fadJ*). Data are mean ± s.d. of biological replicates (n = 3). (D) Medium-chain fatty acid production in the presence of the Fab I inhibitor, triclosan. A. Growth of strain **12** was monitored by OD_600_ growing in the varying concentration of triclosan over 72 h. At 72 h, samples were collected and quantified for octanoic acid production. The octanoic acid productivity of the PaaJ pathway remains unchanged upon addition of triclosan to toxic levels where growth is inhibited. Data are mean ± s.d. of biological triplicates (n = 3).

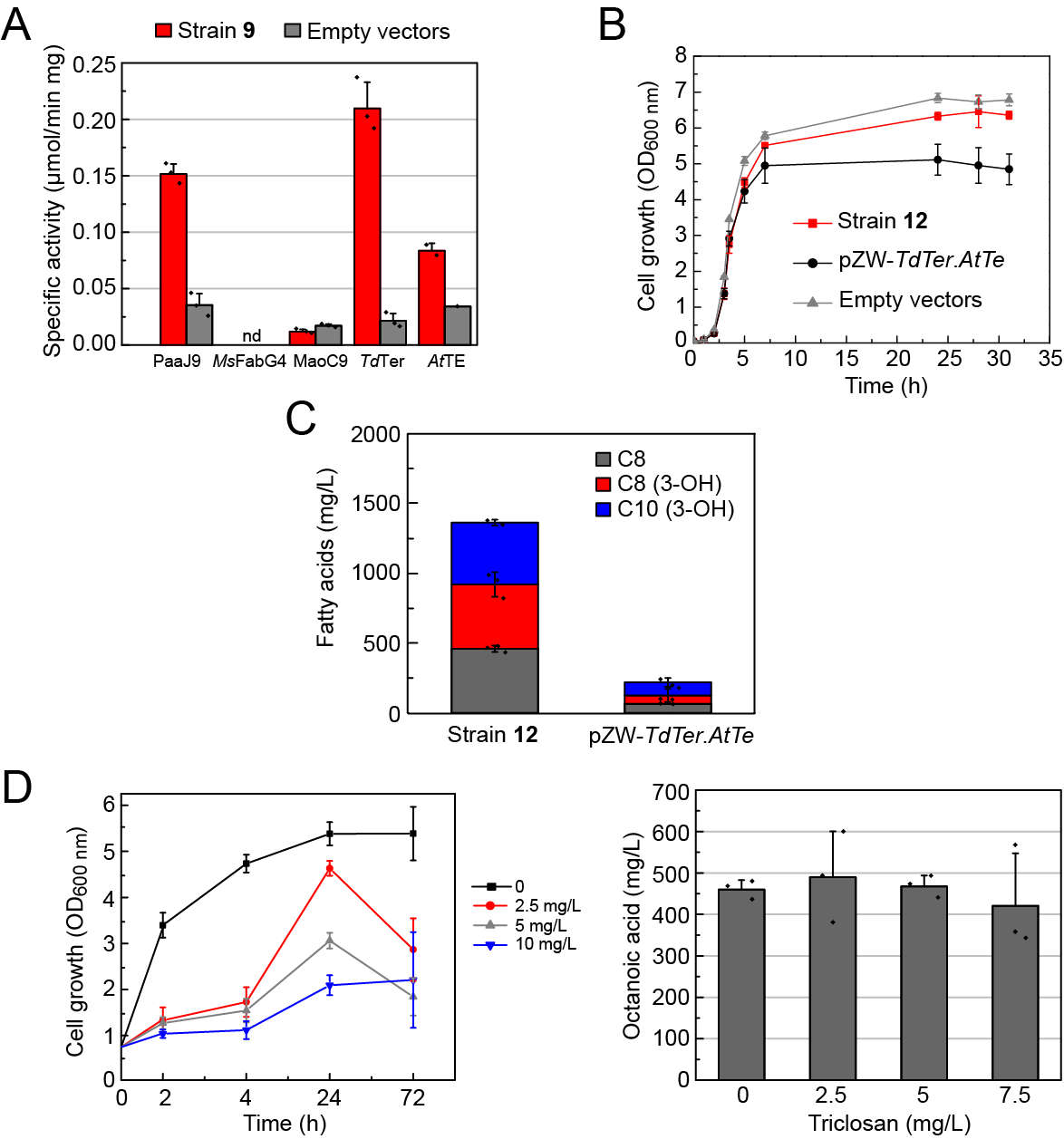

**Figure S8**. Heterogeneous catalyst screening using a 3-hydroxyoctanoic acid standard. 3-Hydroxyoctanoic acid was dissolved in water and passed over commercial catalysts using a microreactor system integrated with a GC-MS at a space velocity of 10 s^-1^. Bar graphs are product selectivities reported at quantitative conversion of substrate (black). The (*n*-1) alkene products (heptenes) are represented in red with the selectivity noted above whereas smaller products from cracking are represented in shades of blue. Larger oligomer products (grey) and CO_2_ (white) are also shown. (A) H-BEA. (B) H-Y. (C) Nb_2_O_5_. (D) ZrO_2_. (E) γ-Al_2_O_3_. (F) Representative mass spectra of the heptenes peak showing isomerization of the initial (*n*-1) terminal alkene.

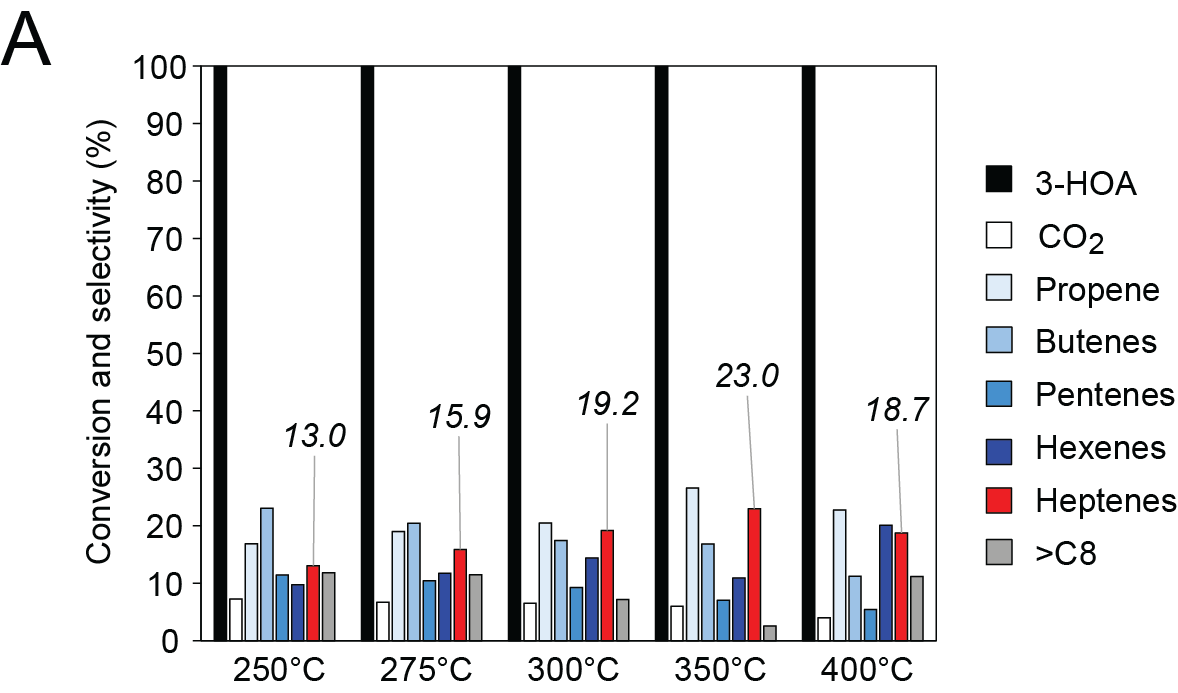

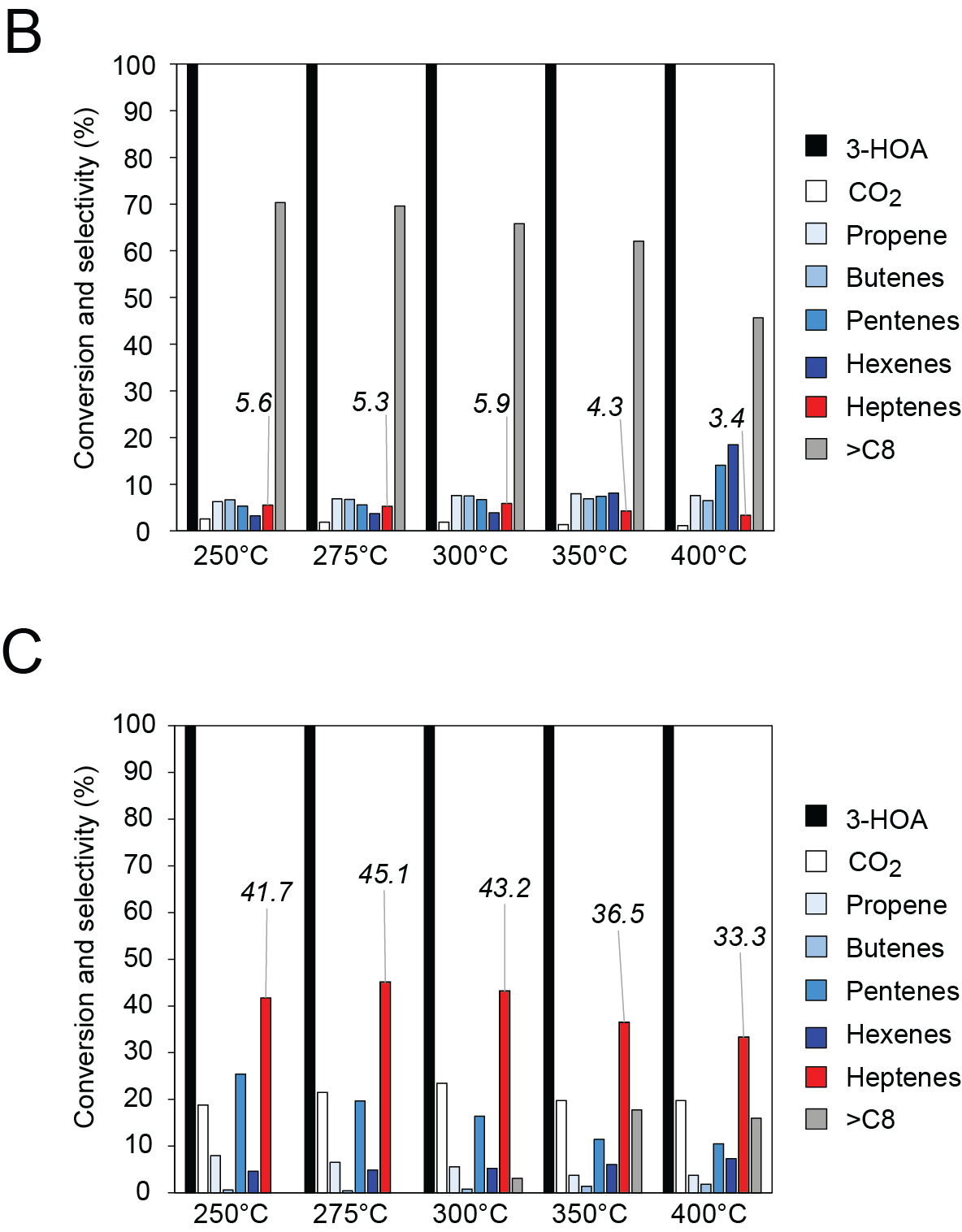

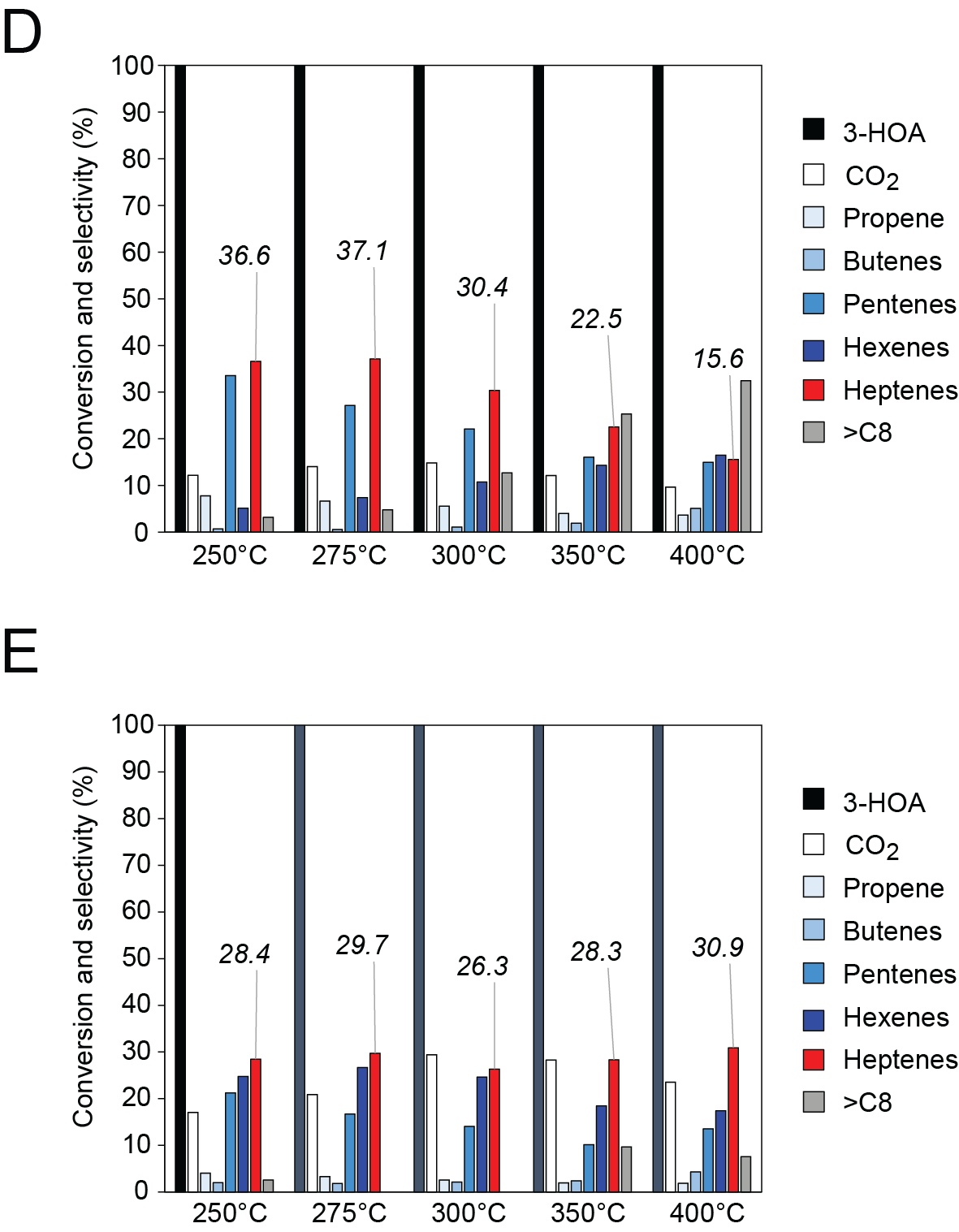

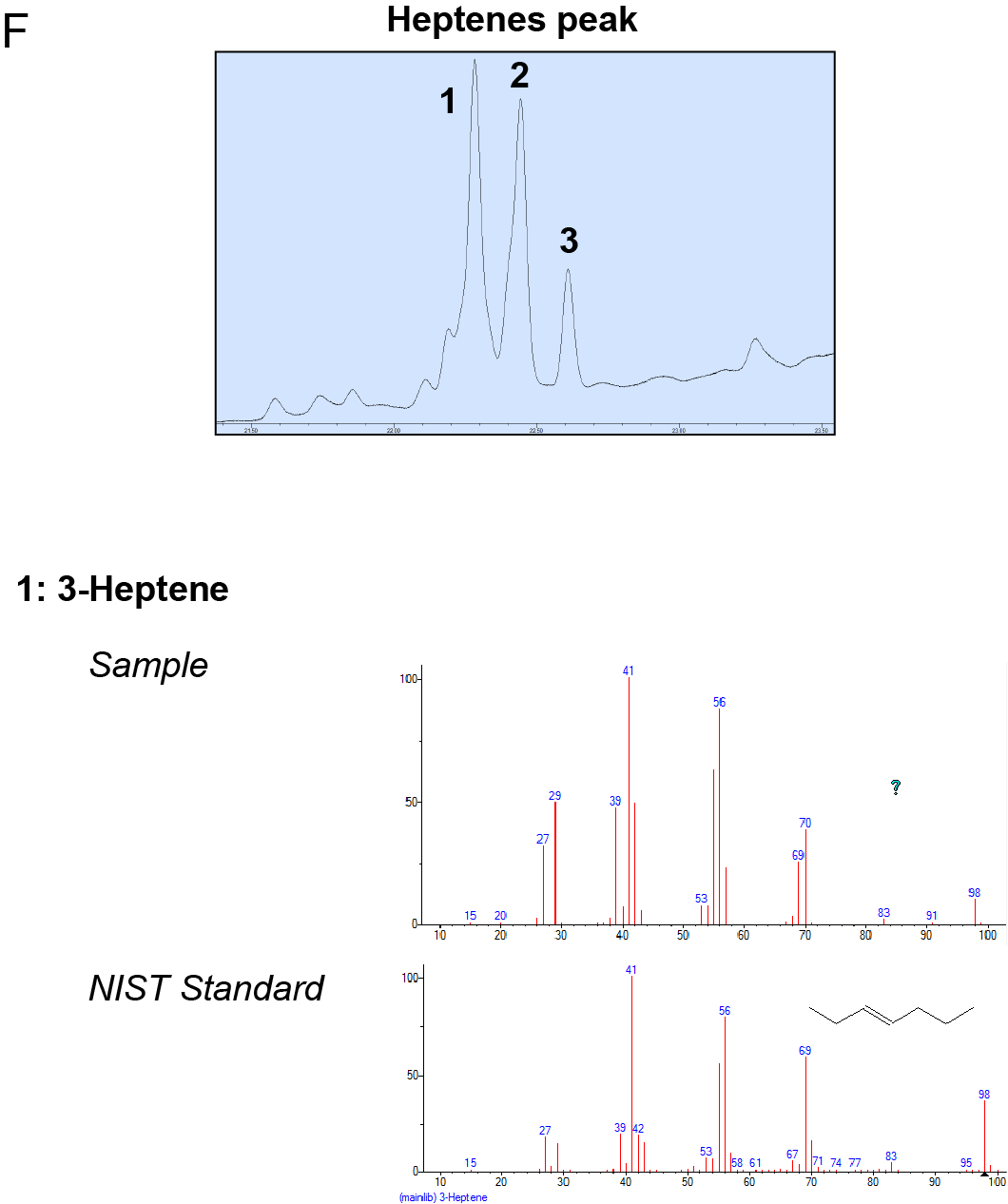

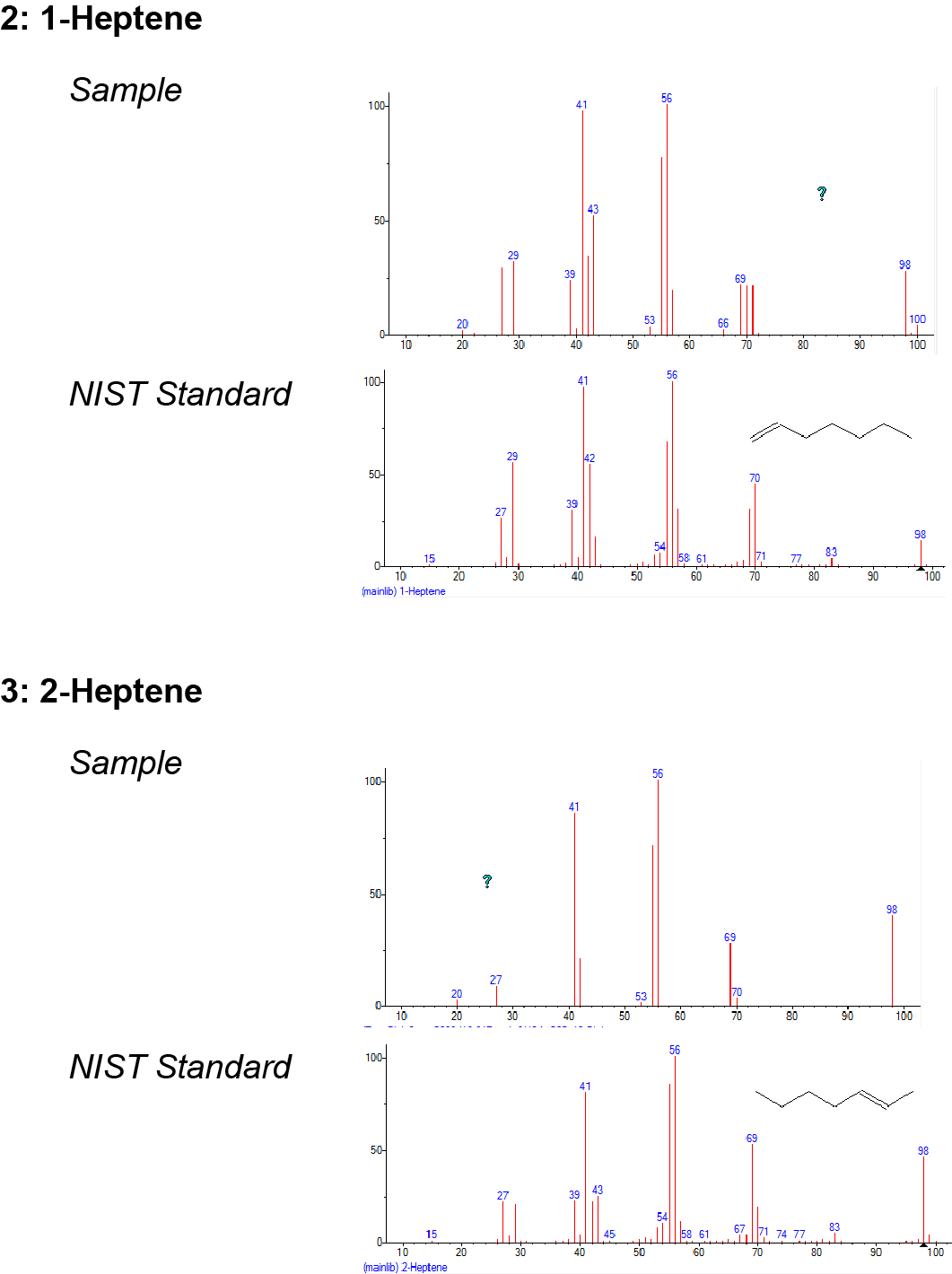
